# Descending neuron population dynamics during odor-evoked and spontaneous limb-dependent behaviors

**DOI:** 10.1101/2022.06.30.497612

**Authors:** Florian Aymanns, Chin-Lin Chen, Pavan Ramdya

## Abstract

Deciphering how the brain regulates motor circuits to control complex behaviors is an important, long-standing challenge in neuroscience. In the fly, *Drosophila melanogaster*, this is accomplished by a population of ∼ 1100 descending neurons (DNs). Activating only a few DNs is known to be sufficient to drive complex behaviors like walking and grooming. However, what additional role the larger population of DNs plays during natural behaviors remains largely unknown. For example, they may modulate core behavioral commands, or comprise parallel pathways that are engaged depending on sensory context. We evaluated these possibilities by recording populations of nearly 100 DNs in individual tethered flies while they generated limb-dependent behaviors. We found that the largest fraction of recorded DNs encode walking while fewer are active during head grooming and resting. A large fraction of walk-encoding DNs encode turning and far fewer weakly encode speed. Although odor context does not determine which behavior-encoding DNs are recruited, a few DNs encode odors rather than behaviors. Lastly, we illustrate how one can identify individual neurons from DN population recordings by analyzing their spatial, functional, and morphological properties. These results set the stage for a comprehensive, population-level understanding of how the brain’s descending signals regulate complex motor behaviors.

## 1 Introduction

The richness of animal behaviors depend on the coordinated actions of many individual neurons within a population. For example, individual neurons or small networks may compete in a winner-take-all manner to select the next most appropriate motor action [1]. The brain can then convey these decisions to motor circuits via a population of descending neurons (DNs) projecting to the spinal cord of vertebrates, or ventral nerve cord (VNC) of invertebrates. There, DNs axons impinge upon local circuits including central pattern generators (CPGs) that transform DN directives into specific limb or body part movements [2–5]. DNs make up only about 1 % of brain neurons. Thus, DN population activity represents a critical information bottleneck: high-dimensional brain dynamics must be compressed into low-dimensional commands that efficiently interface with and can be read out by motor circuits. The information carried by individual DNs has long been a topic of interest [6– 8]. Through electrophysiological recordings in large insects, the activities of individual DNs have been linked to behaviors like walking and stridulation [6]. For some DNs links between firing rate and behavioral features like walking speed have also been established [8]. However, how the larger population of DNs coordinate their activities remains unknown.

The fruit fly, *Drosophila melanogaster*, is an excellent model for investigating how DNs regulate behavior. Flies are genetically-tractable, have a rich behavioral repertoire, and also have a numerically small and compact nervous system [9]. There are only thought to between ∼ 350 [10] and ∼ 550 [11] pairs of DNs in *Drosophila*. Sparse sets of these DNs can be experimentally targeted using transgenic driver lines [10] for functional recordings [12–16], or optogenetic perturbations [16–20]. Their functional properties can be understood within a circuit context using emerging connectomics datasets [21, 22]. Thus, building upon foundational work in other insects [6–8], studies in *Drosophila* can ultimately reveal how identified DNs work collectively to regulate complex behaviors.

Until now, investigations of *Drosophila* have focused on individual or small sets of of DNs. These studies have demonstrated that artificial activation of DN pairs is sufficient to drive complex behaviors including escape [23](giant fiber neurons, GF), antennal grooming [24](antennal Descending Neurons, aDN), backward walking [25](Moonwalker Descending Neurons, MDN), forward walking [26](DNp09), and landing [19](DNp10 and DNp07). These results also suggest a command-like role for some DNs, in that they are both necessary and sufficient to drive particular actions [27].

Although a command-like role for individual DNs is intuitively easy to grasp, it may not translate well toward understanding how natural behavior is coordinated by large DN populations. Notably, a behavioral screen revealed that optogenetic activation of most DNs drives only small changes in locomotor and grooming behaviors [17], rather than a large variety of distinct actions as might be expected if each DN was a command-like neuron. Therefore, DN populations are likely employing additional, alternative control approaches. For example, some groups of DNs might modulate or fine-tune actions primarily driven by other, command-like DNs. The balance between command-like and modulatory roles of DNs may differ for stereotyped versus more flexible behaviors. In line with their role in behavioral modulation, studies in crickets and locusts have demonstrated that changes in the firing rates of some DNs [8, 28] correlate with walking speed and turning [6]. Alternatively, DN subpopulations may represent parallel pathways that are recruited depending on sensory context. For example, different groups of DNs may be differently engaged during odor-evoked versus spontaneously-generated walking [6,7]. Finally, some DNs may convey raw sensory information to enable feedback control by downstream motor circuits. For example, in addition to discovering DNs active during steering, one recent study also observed DNa02 activity in response to fictive odors in immobile flies [29]. DNp07 activity has also been observed in non-flying flies in response to visual stimulation [19].

To resolve the degree to which DNs (i) modulate ongoing behaviors, (ii) are recruited depending on sensory context, and/or (iii) convey raw sensory information from the environment, one would need to simultaneously record large numbers of DNs in a behaving animal. Until now, this has been technically difficult for several reasons. First, there has been an absence of tools for selectively genetically targeting DN populations. Additionally, because DN cell bodies and neurites are distributed across the brain [10], relatively invasive [30] volumetric imaging would be required to simultaneously record the activity of many at once. We previously developed a thoracic dissection approach that enables the optical recording of DN axons within the cervical connective in tethered, behaving animals [14, 31]**(Figure 1a)**. Here, we combined this imaging approach with genetic tools [32] that restrict the expression of neural activity reporters to the brain to record populations of nearly 100 DNs in individual tethered, behaving flies. During these recordings, we presented olfactory stimuli and acquired behavioral data for 3D pose estimation [33], as well as fictive locomotor trajectories [34].

**Figure 1:**
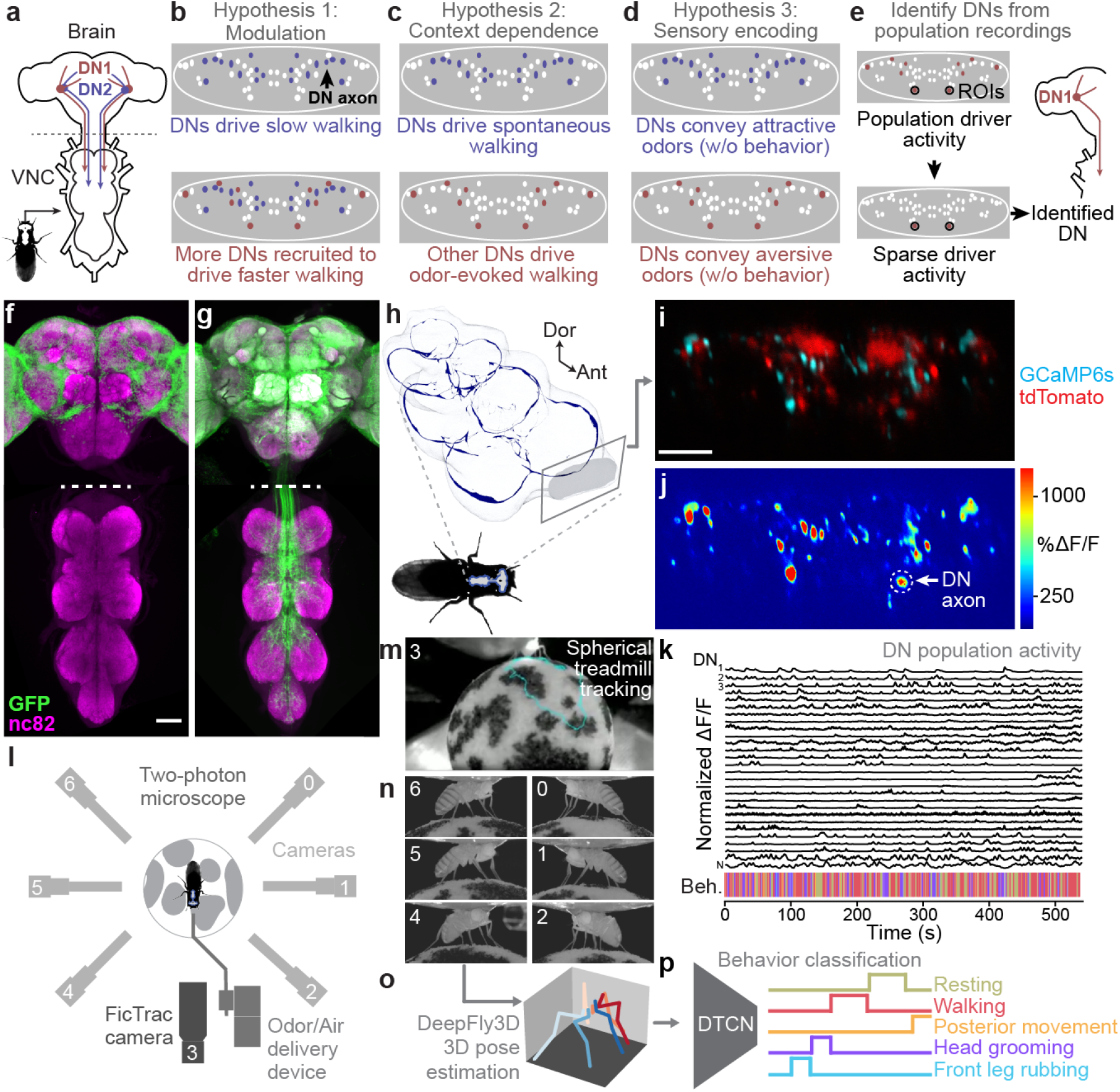
Recording descending neuron population activity and animal behavior. **(a)** Schematic of the *Drosophila* nervous system showing descending neurons (DNs) projecting from the brain to motor circuits in the ventral nerve cord (VNC). Only two pairs of DNs (red and blue) are shown for clarity. Indicated (dashed gray line) is the coronal imaging region-of-interest in the thoracic cervical connective. **(b)** In a ‘modulation’ framework for DN population control, new DNs (red) may be recruited to modulate ongoing behaviors primarily driven by core DNs (blue). Each ellipse is an individual DN axon (white, blue, and red). **(c)** In a ‘context dependence’ framework for DN population control, different DNs may be recruited to drive identical behaviors depending on sensory context. **(d)** Alternatively, in a ‘sensory encoding’ framework, many DNs may not drive behaviors but rather transmit sensory signals to the VNC. **(e)** An approach for identifying DNs from population recordings. One may first identify sparse transgenic strains labeling specific neurons from DN populations (circled in black) using their functional attributes/encoding, positions within the cervical connective, and axon shapes. Then, one can use sparse morphological data to find corresponding neurons in the brain and VNC connectome. **(f**,**g)** Template-registered confocal volume z-projections illustrating a ‘brain only’ driver line (otd-nls:FLPo; R57C10-GAL4,tub*>*GAL80*>*) expressing **(f)** a nuclear (histone-sfGFP) or **(g)** a cytosolic (smGFP) reporter. Scale bar is 50 µm. Location of twophoton imaging plane in the thoracic cervical connective is indicated (white dashed lines). Tissues are stained for GFP (green) and neuropil (‘nc82’, magenta). **(h)** Schematic of the VNC illustrating the coronal (x-z) imaging plane. Dorsal-ventral (‘Dor’) and anterior-posterior (‘Ant’) axes are indicated. **(i)** Denoised two-photon image of descending neuron (DN) axons passing through the thoracic cervical connective. Scale bar is 10 µm. **(j)** Two-photon imaging data from panel **i** following motion correction and %Δ*F/F* color-coding. An ROI (putative DN axon) is indicated (white dashed circle). **(k)** Sample normalized %Δ*F/F* time-series traces for 28 (out of 95 total) DN axons recorded from one animal. Behavioral classification at each time-point is indicated below, color-coded is as in panel **p. (l)** Schematic of system for recording behavior and delivering odors during two-photon imaging (not to scale) while a tethered fly walks on a spherical treadmill. **(m)** Spherical treadmill ball rotations (fictive walking trajectories) are captured using the front camera and processed using FicTrac software. Overlaid (cyan) is a sample walking trajectory. **(n)** Video recording of a fly from six camera angles. **(o)** Multiview camera images are processed using DeepFly3D to calculate 2D poses and then triangulated 3D poses. These 3D poses are further processed to obtain joint angles. **(p)** Joint angles are input to a dilated Temporal Convolutional Network (DTCN) to classify behaviors including walking, resting, head (eye and antennal) grooming, front leg rubbing, or posterior (abdominal and hindleg) movements.

Using these tools, we could test the extent to which DN population activity patterns are consistent with roles in behavior modulation **(Figure 1b)**, context-dependence recruitment **(Figure 1c)**, and/or raw sensory signaling **(Figure 1d)**. We observed that the largest fraction of DNs are active during walking. Principal component analysis revealed diverse neural dynamics across epochs of walking. This variability reflected partially overlapping subsets of DNs that encode turning and, to a far weaker extent, speed. These data support a role for DN populations in behavioral modulation. DNs are active during walking or grooming irrespective of whether the behavior was generated spontaneously or during olfactory stimulation. These data rule out a strong context dependence for the recruitment of DN subpopulations. However, we did find that some DNs are specifically responsive to odors, revealing that motor circuits have access to surprisingly unfiltered sensory information. Finally, we illustrate how one can identify DNs from population recordings **(Figure 1e)**. We studied a prominent pair of DNs that are asymmetrically active during antennal grooming. By using their topological and encoding properties, we could identify a sparse driver line that targets these neurons used morphological analysis (MCFO and connectomics) to confirm their identities as DNx01 neurons originating from the antennae [10]. These data provide a first, global view of DN population activity during natural behaviors and open the door to a comprehensive mechanistic understanding of how the brain’s descending signals regulate motor control.

## 2 Results

### 2.1 Recording descending neuron population activity in tethered, behaving *Drosophila*

To selectively record the activity of populations of DNs, we devised an intersectional genetic and optical approach. First, we restricted the expression of GCaMP6s (a fluorescent indicator of neural activity [35]) and tdTomato (an anatomical fiduciary) to the supraesophageal zone of the brain [32](*;otdnls:FLPo* ; *R57C10-GAL4, tub>GAL80>*). We confirmed that transgene expression was restricted to cell bodies in the brain **(Figure 1f)**. The axons of DNs could be seen passing through the cervical connective, targeting motor circuits within the ventral nerve cord (VNC) **(Figure 1g)**. Thus, although our driver line lacks expression in the subesophageal zone (SEZ) **(Figure S1c)**—a brain region known to house at least 41 DNs [10, 36] some of which can drive grooming [20](DNg11, DNg12)— we could still capture the activities of a large population of DNs. Second, by performing coronal (x-z) two-photon imaging of the thoracic cervical connective [14], we could exclusively record DN axons **(Figure 1h)** in tethered animals that were behaving on a spherical treadmill. This imaging approach could also compensate for image translations during animal behavior, keeping regions-of-interest (ROIs) within the field-of-view (FOV). We then applied image registration [14]—to correct for translations and deformations—as well as image denoising [37]—to obtain higher signal-to-noise images. We confirmed that denoising does not systematically delay or prolong the temporal dynamics of neural activity **(Figure S1a,b)**. Resulting images clearly showed elliptical ROIs that likely represent individual large axons or possibly also tightly packed groups of smaller axons **(Figure 1i)**. From here on we will interchangeably refer to ROIs as DNs or neurons. From these data, we calculated Δ*F/F* images **(Figure 1j)** within which we could manually label 75-95 of the most distinct and clearly visible ROIs in each animal. This resulted in high-dimensional neural time-series **(Figure 1k)**.

To test the context-dependence and sensory feedback encoding of DNs, we built an olfactometer that could sequentially present humidified air, and one of two alternating odors: apple cider vinegar (‘ACV’, an attractive odorant [38]), or methyl salicylate (‘MSC’, a putatively aversive odorant [39]). During experiments, we alternated presentation of ACV and MSC with humid air. We performed photoionization detector (PID) measurements to confirm that our olfactometer could deliver a steady flow of air/odor with minimal mechanical perturbations during switching **(Figure S1d-f)**.

Along with neural recordings and odor delivery, we quantified limb and joint positions by recording tethered animals from six camera angles synchronously at 100 frames-per-second (fps) **(Figure 1l)**. A seventh, front-facing camera recorded spherical treadmill rotations that could be converted into fictive locomotor trajectories using FicTrac [34] **(Figure 1m)**. Multi-view camera images **(Figure 1n)** were post-processed using DeepFly3D [33] to estimate 3D joint positions [40] **(Figure 1o)**. These data were used to train a dilated temporal convolutional neural network (DTCN) [41] that could accurately classify epochs of walking, resting, head (eye and antennal) grooming, front leg rubbing, and posterior (hindleg and abdominal) movements **(Figure 1p, Figure S1g)**. Animals predominantly alternated between resting, walking, and head grooming with little time spent front leg rubbing or moving their posterior limbs and abdomen **(Figure S1h)**. Notably, we also observed structure in our behavioral data: flies were more likely to walk after resting or generating posterior movements **(Figure S1i)**. Flies also frequently performed front leg rubbing following head grooming [42] **(Figure S1i)**. Taken together, this experimental and computational pipeline yielded a rich dataset of DN population activity and associated odor-evoked and spontaneous behaviors **(Video 1)**.

### 2.2 Encoding of behavior in descending neuron populations

With these data, we first asked to what extent DN populations encode—and presumably regulate— each of our classified behaviors: resting, walking, head grooming, posterior movements, and front leg rubbing. We addressed this question in two ways. First, we asked how well each behavior could be predicted based on the activity of each neuron. Specifically, we used a linear model to quantify the extent to which a given DN’s activity could explain the variance of (i.e., ‘encode’) each behavior **(Figure 2a,b)**. The largest fraction (∼60%) of DNs encode walking. The second largest group of DNs encode head grooming (∼15%). Only a very small fraction of DNs encode resting and no neurons encode front leg rubbing, or posterior movements **(Figure 2c)**. However, some of these behaviors were very infrequent (posterior movements) or of short duration (front leg rubbing) **(Figure S1h)**, weakening the power of our analysis in these cases. As well, although none of the DNs *best* explained posterior movements and front leg rubbing out of all behaviors, we observed that DNs encoding walking also encoded posterior movements. Similarly, DNs encoding head grooming also encoded front leg rubbing **(Figure 2b)**. This may be due to the strong sequential occurrence of these pairs of behaviors **(Figure S1i)** and the long decay time constant of GCaMP6s (∼1 s [35]) resulting in elevated calcium signals for the subsequent behavior in the pair. To resolve the extent to which these DNs truly encode one behavior versus the other, we performed a more narrow linear regression analysis. We used equal amounts of data from this pair of sequential behaviors, and calculated the neural variance (rather than the behavioral variance) uniquely explained by these two behaviors alone. These analyses confirmed that DNs predominantly encode walking rather than posterior movements **(Figure S2a)**, and head grooming rather than front leg rubbing **(Figure S2b)**.

**Figure 2:**
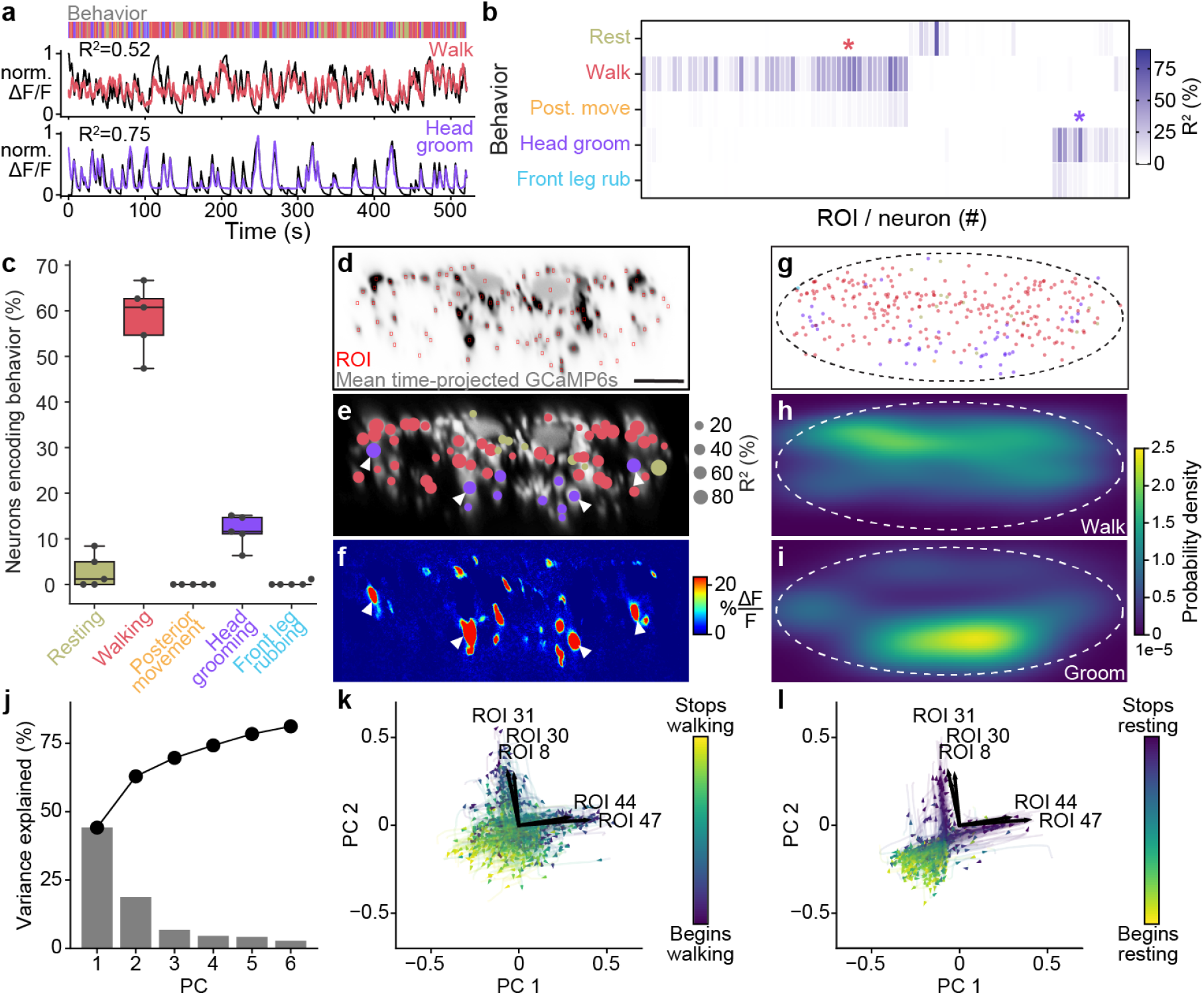
Encoding of behavior in descending neuron populations. **(a)** Shown for walking (top) and head grooming (bottom) are the activity (normalized and cross-validation predicted Δ*F/F*) of individual walk-and head-groom encoding DNs (red and purple lines), as well as predicted Δ*F/F* traces from convolving binary behavior regressors with a crf (black lines). The output of the behavior classifier is shown (color bar). **(b)** The cross-validation mean of behavioral variance explained by each of 95 DNs from one animal. Colored asterisks are above the two DNs illustrated in panel **a. (C)** The percentage of DNs encoding each classified behavior across five animals. Box plots indicate the median, lower, and upper quartiles. Whiskers signify 1.5 times the interquartile range. **(d)** Mean time projection of GCaMP6s fluorescence over one nine minute recording. Image is inverted for clarity (high mean fluorescence is black). Manually identified DN regions-of-interest (ROIs) are shown (red rectangles). Scale bar is 10 µm. Panels **d-i** share the same scale. **(e)** DNs color-coded (as in panel **c**) by the behavior their activities best explain. Radius scales with the amount of variance explained. Prominent head groom-encoding neurons that are easily identified across animals are indicated (white arrowheads). **(f)** Behavior-triggered average Δ*F/F* image for head grooming. Prominent head grooming DNs identified through linear regression in panel **e** are indicated (white arrowheads). **(g)** Locations of DNs color-coded by the behavior they encode best. Data are from five animals. **(h**,**i)** Kernel density estimate based on the locations of **(h)** walking or **(i)** head grooming DNs in panel **g. (j)** Amount of variance explained by the principal components (PCs) of neural activity derivatives during walking. **(k**,**l)** Neural activity data during **(k)** walking and **(l)** resting evolve on two lobes. The PC embedding was trained on data taken during walking only. Colored lines indicate individual epochs of **(k)** walking and **(l)** resting. Time is color coded and the temporal progressions of each epoch is indicated (arrowheads). Note that color scales are inverted to match the color at transitions between walking and resting. Black arrows indicate ROIs with high PC loadings. ROI number corresponds to the matrix position in panel **b**. For their locations within this fly’s cervical connective, see **Figure S5d**.

We next asked to what extent DNs encode—and presumably drive—the kinematics of joints, limbs, or limb pairs rather than behaviors. To test this possibility, we quantified how much better neural activity could be predicted from joint angles rather than from behavior. Specifically, we computed the amount of variance in DN activity that could be uniquely explained by subgroups of joint angles, but not behavior categories or any of the remaining joint angles. Separating joint angles into groups (all, pairs, or individual legs) allowed us to probe the possibility that some neurons might control pairs of legs or individual legs and helped to mitigate the effect of correlations between joint angles within individual legs. Because of the rapid movements of each leg (5–20 Hz [43, 44]) and the long decay time of our calcium indicator, we also convolved joint angle and behavior regressors with a calcium response function (crf) kernel. We found that joint angles can only very marginally improve the prediction of neural activity beyond simply using behavior regressors **(Figure S4)**. Thus, DN populations largely encode high-level behaviors, suggesting that they delegate low-level kinematic control to downstream circuits in the VNC.

### 2.3 The spatial organization of descending neuron encoding

Previous morphological analyses demonstrated a clear organization of *Drosophila* DN projections within the VNC [10]. This is likely linked to the distinct functional partners of DNs which regulate limb-dependent (e.g., walking and grooming) versus wing-dependent (e.g., flight and courtship display) behaviors. To further explore the relationship between function and topology, we next asked to what extent we might observe a relationship between a DN’s encoding of behavior and its axon’s position within the cervical connective **(Figure 2d)**. We found that DNs encoding walking are spread throughout the dorsal connective **(Figure 2e,g,h)**. On the other hand, head groom-encoding DNs are predominantly in the ventral connective **(Figure 2g,i)** including two prominent pairs—lateral and medial-ventral **(Figure 2e, white arrowheads)**—whose activities explain the largest amount of variance in head grooming across animals **(Figure S3c)**. Surprisingly, we also observed DNs which encode resting. These were located medially, close to the giant fibers, as well as in the lateral extremities of the connective **(Figure 2e, olive circles)**. We speculate that rest-encoding DNs may suppress other behaviors or could actively drive the tonic muscle tone required to maintain a natural posture.

We next performed a complementary analysis to further examine the functional-topological organization of DNs in the connective. Specifically, we generated behavior-triggered averages of Δ*F/F* images. The results of this approach were only interpretable for frequently occurring behaviors so here we focused on walking, resting, and head grooming **(Figure S1h)(Videos 2-4)**. Across animals, we consistently observed DNs encoding walking in the dorsal connective, and two pairs of ventral DNs encoding head grooming **(Figure 2f; Figure S3d)**. By contrast, rest-encoding DNs were located in less consistent locations within the connective. These findings confirm that DN populations for walking and head grooming are largely spatially segregated, a feature that may facilitate the identification of specific cells from DN population recordings across animals.

### 2.4 Descending neuron population dynamics suggest more nuanced feature encoding

Thus far we have observed that DN subpopulations encode distinct behaviors and that, by far, the largest fraction encode walking. However, locomotion is not monolithic but rather continuously varies in speed and direction within a single walking trajectory. Thus, the large number of DNs active during walking may represent an aggregate of subpopulations that are differently engaged during distinct locomotor modes. To address this hypothesis, we first closely examined the temporal structure of DN population activity dynamics only during walking epochs. We asked to what extent there is variability and structure in population activity that could potentially support the modulatory encoding of walking speed, forward/backward, or turning.

As in a similar analysis of *C. elegans* population dynamics [45], we calculated the temporal derivative of each DN’s activity and then performed principal component analysis (PCA) on these time-series. We found that the first two PCs can explain upwards of 60 % of the variance in DN population activity during walking **(Figure 2j; Figure S5a)**. Visualizing 2D trajectories (PC 1 and PC 2 subspace) of DN activity during individual walking bouts revealed that it moved primarily along two directions **(Figure 2k; Figure S5b)**. These directions were even more clear for resting data (embedded within the same PC space) just before the fly began to walk **(Figure 2l; Figure S5c)**. To identify individual DNs that most heavily influence this dynamical divergence, we next found those with the largest PC loadings. These neurons’ activities most strongly influence the position of population activity in the PC space **(Figure 2k,l; Figure S5b,c)**. Consistently, also in flies with a less clear divergence in neural trajectories, we found subsets of DNs whose activities correspond to one of these two directions. By examining the positions of their axons, we observed that they are spatially segregated on opposite sides of the connective **(Figure S5d, cyan arrowheads)**.

### 2.5 Descending neurons that encode walking include spatially segregated turn-encoding clusters

The divergence of population dynamics and spatial segregation of associated neurons led us to hypothesize that subsets of walk-encoding DNs might be preferentially become active during left and right turning. Alternatively, they might encode fast versus slow walking speeds [26]. Studies in other insects and vertebrates have shown that DNs can play a modulatory role by regulating turning and speed during locomotion [3, 6]. As well, in *Drosophila*, the activation of DNp09 neurons can drive forward walking [26]. Recordings from sparse sets of DNa01 [14, 29] and DNa02 [29] neurons have also shown turn encoding (i.e., steering).

Therefore, we next tested if variability in the activity of DNs might reflect fine-grained encoding of turning and speed during forward walking. We did not analyze backward walking due to its scarcity **(Figure S6a)**, brevity **(Figure S6b)** and minimal dynamic range **(Figure S6c)** in our experimental data. Specifically, we quantified the degree to which DN population activity could uniquely explain yaw (turning) or pitch (speed) angular velocity of the spherical treadmill. Both of these time-series data were convolved with a crf to account for slow calcium indicator decay dynamics. To capture information about turning and walking speed that could not simply be explained by whether the fly was walking or not, we compared the explained variance of our neuron-based ridge regression to a model predicting these features from just a binary walking regressor and shuffled neural data **(Figure 3a)**. Neural activity could explain a great deal of variance in turning and, to a lesser extent, speed. Here, the absence of speed encoding may be because the binary walking regressor alone can partially predict speed variance from transitions between resting and walking (*R*^2^ = 25 %). Therefore, we refined our analysis by only using data from walking epochs and calculating the amount of turning and speed variance that could be explained using neural activity. In this manner, we confirmed that neural activity can uniquely explain turning to a greater extent (∼ 60 %) than it can explain walking speed (∼ 30 %) (**Figure 3b**).

**Figure 3:**
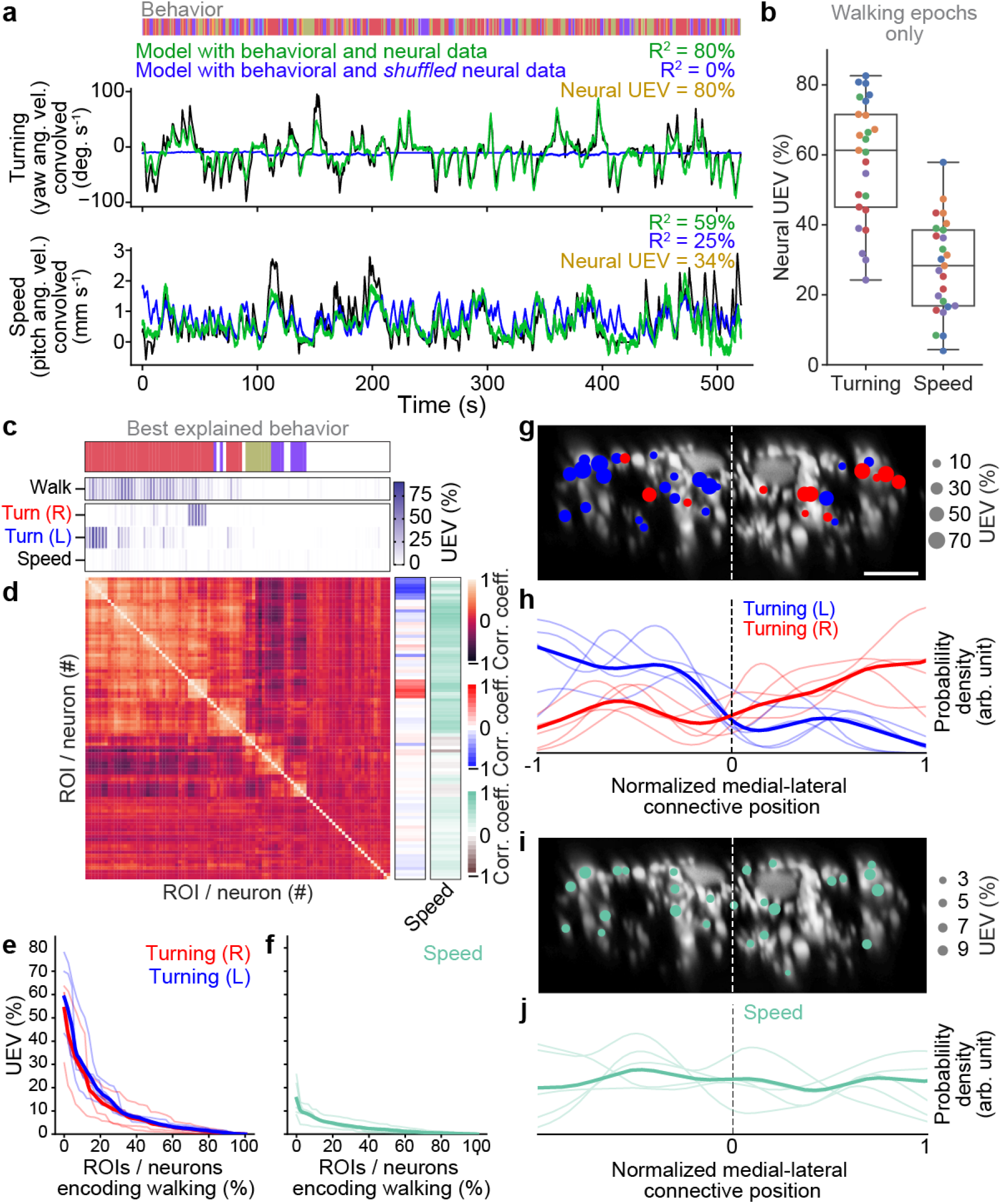
Turning and speed encoding in descending neuron populations. **(a)** Predictions of (top) turning and (bottom) walking speed modeled using convolved behavior regressors and all neurons in one animal. Shown are predictions (green) with all regressors intact or (blue) with neural data shuffled across time. Indicated are *R*^2^ values obtained by comparing predicted and real (black) turning and walking speed. These are subtracted to obtain neural unique explained variance (UEV). The fly’s behavior throughout the recording is indicated (color bar). **(b)** UEV obtained only using data taken during walking, thus accounting for trivial explanations of speed and turning variance resulting from transitions between resting and walking. Shown are data from five trials each for five flies (color-coded). **(c)** UEV of each DN from one animal for walking speed or left and right turning ordered by clustering of Pearson’s correlation coefficients in panel **d**. Walking *R*^2^ values are the same as in **Figure 2b** but reordered according to clustering. The models for turning and walking speed were obtained using behavior regressors as well as neural activity. To compute the UEV, activity for a given neuron was shuffled temporally. The behavior whose variance is best explained by a given neuron is indicated (color bar). **(d)** Pearson’s correlation coefficient matrix comparing neural activity across DNs ordered by clustering. Shown as well are the correlation of each DN’s activity with right, left, and forward walking (right). **(e**,**f)** UEV for **(e)** turning or **(f)** speed for DNs that best encode walking. Neurons are sorted by UEV. Shown are the distributions for individuals (translucent lines), and the mean across all animals (opaque line). **(g)** Locations of turn-encoding DNs (UEV *>* 5%), color-coded by preferred direction (left, blue; right, red). Circle radii scale with UEV. Dashed white line indicates the approximate midline of the cervical connective. Scale bar is 10 µm for panels **g** and **i. (h)** Kernel density estimate of the distribution of turn encoding DNs. Shown are the distributions for individuals (translucent lines), and the mean distribution across all animals (opaque lines). Probability densities are normalized by the number of DNs along the connective’s medial-lateral axis. **(i)** Locations of speed encoding DNs (UEV *>* 2%). Circle radii scale with UEV. Dashed white line indicates the approximate midline of the cervical connective. **(j)** Kernel density estimate of the distribution of speed encoding DNs. Shown are the distributions for individuals (translucent lines), and the mean distributions across all animals (opaque lines). Probability densities are normalized by the number of DNs along the connective’s medial-lateral axis.

We next investigated which individual or groups of neurons contribute to the prediction of turning and walking speed. To do this, we only used the activity of one neuron at a time in our model (by contrast, in the paragraph above we used all neurons). Among DNs that encode walking, we found specific DNs that strongly explain right or left turning. By contrast, a more distributed set of DNs weakly encode walking speed **(Figure 3c; Figure S7a)**. Having identified clusters of DNs encoding right (‘red’) and left (‘blue’) turning, we next investigated whether there might be groups encoding other walking features. Among neurons that best explain walking, we again observed clusters for turning but no prominent clusters for speed **(Figure 3d; Figure S7b)**. This was also reflected in the amount of variance explained: across animals, some walk-encoding DNs also strongly encoded turning **(Figure 3e)**, whereas these DNs only encoded a tiny fraction of the variance in walking speed **(Figure 3f)**.

Simple models for locomotor control [46] suggest that turning can be controlled by the relative activities of DNs on one side of the brain versus the other. This is supported by a study showing asymmetric activation of *Drosophila* MDNs [47]. Alternatively, the spatial asymmetry in neural activity required for turning might arise in circuits downstream of DNs within the VNC (i.e., with no spatial asymmetry in DN activity). To distinguish between these possibilities, we examined the spatial location of turn-encoding DNs in the cervical connective. We found both ipsi- and contralateral turn-encoding DNs on both sides of the connective **(Figure 3g; Figure S7c)**, but a clear ipsilateral enrichment **(Figure 3h; Figure S7d)**. Many of the DNs encoding turning had high PC loading **(Figure S5d)** revealing that turning contributes heavily to the variance in DN population dynamics during walking. By contrast, DNs encoding walking speed were more homogeneously distributed across the connective with no clear spatial enrichment **(Figure 3i,j; Figure S7e,f)**. Overall, these data support the notion that, during walking, DN population activity largely varies due to turn-related modulation rather than shifts in walking speed.

### 2.6 Descending neurons are active during behaviors irrespective of olfactory context

Beyond a modulatory role, the large number of DNs active during walking could reflect context dependence: specific subpopulations may only be engaged as a function of sensory context. The possibility of recruiting separate pools of DNs for walking is supported by the observation that an attractive odor, apple cider vinegar (ACV), decreases resting and increases walking **(Figure S8a)**, increases forward walking speed **(Figure S8b)**, and reduces turning **(Figure S8c,d)**.

To address the extent to which subgroups of DNs are recruited depending on olfactory context, we studied the amount of walking or head grooming variance explained by each DN using only data acquired during exposure to either humidified air, apple cider vinegar (ACV), or methyl salicylate (MSC)—rather than analyzing all walking epochs as in our previous analysis. Humidified air data were subsampled to match the smaller amount of data available for ACV and MSC presentation. We found that largely the same DNs were highly predictive of walking **(Figure S9a)** and head grooming **(Figure S9b)** irrespective of olfactory context. Only a very small fraction of DNs were differentially recruited during odor presentation **(Figure S9a,b, black asterisks)**. Of these four DNs with different recruitment, three achieve significance because they have only a few values distinct from zero in a single trial. The overall explained variance is also very small **(Figure S9c)**. These data suggest that changing odor context alters action selection and locomotor kinematics but does not shift the identity of active DN subpopulations driving behavior.

### 2.7 Descending neurons exhibit raw odor encoding

Although walk- and head groom-encoding DN populations are recruited irrespective of odor context, it has been shown that a fictive odor (i.e., optogenetic activation of *Orco>CsChrimson*) can activate DNa02 neurons in immobile animals [29]. This implies that DNs may encode the presence and identity of real odors. To examine the extent of this raw sensory encoding, we trained and cross-validated linear discriminant odor classifiers using neural residuals (i.e., the neural activity remaining after subtracting activity that could be predicted using a model based on crf convolved behavior regressors). These residuals allowed us to control for the fact that odor presentation also modulates behavioral statistics **(Figure S8a-c)**.

Classification using neural residuals performed significantly better than classification using behavioral information alone (*p <* 0.0001 for a two-sided Mann-Whitney U test)**(Figure 4a)**. This reveals raw odor encoding within DN populations. However, this might result from many neurons with weak, complementary odor encoding or a few neurons with strong odor encoding. To distinguish between these possibilities, we next identified which DNs encode olfactory signals. We predicted each neuron’s activity using regressors for behavior and the presence of each odor. The more intuitive approach of modeling neural activity by just using odor regressors does not account for the confound that behaviors are also modulated by specific odors. Therefore, we computed the amount of neural variance that could uniquely be explained by the presence of an odor and none of the behavior variables. We found that the activity of a few DNs could be uniquely explained by each of the odors **(Figure 4b, asterisks)**. In these neurons, clear activity peaks coincided with the presence of MSC **(Figure 4c)** or ACV **(Figure 4d)**. Notably, there appears to be no overlap between the DNs encoding MSC or ACV within individual animals **(Figure S8e)**. MSC encoding neurons were found dorsally on the lateral sides of the giant fibers in the connective while ACV encoding neurons were more broadly dispersed **(Figure 4e,f)**.

**Figure 4:**
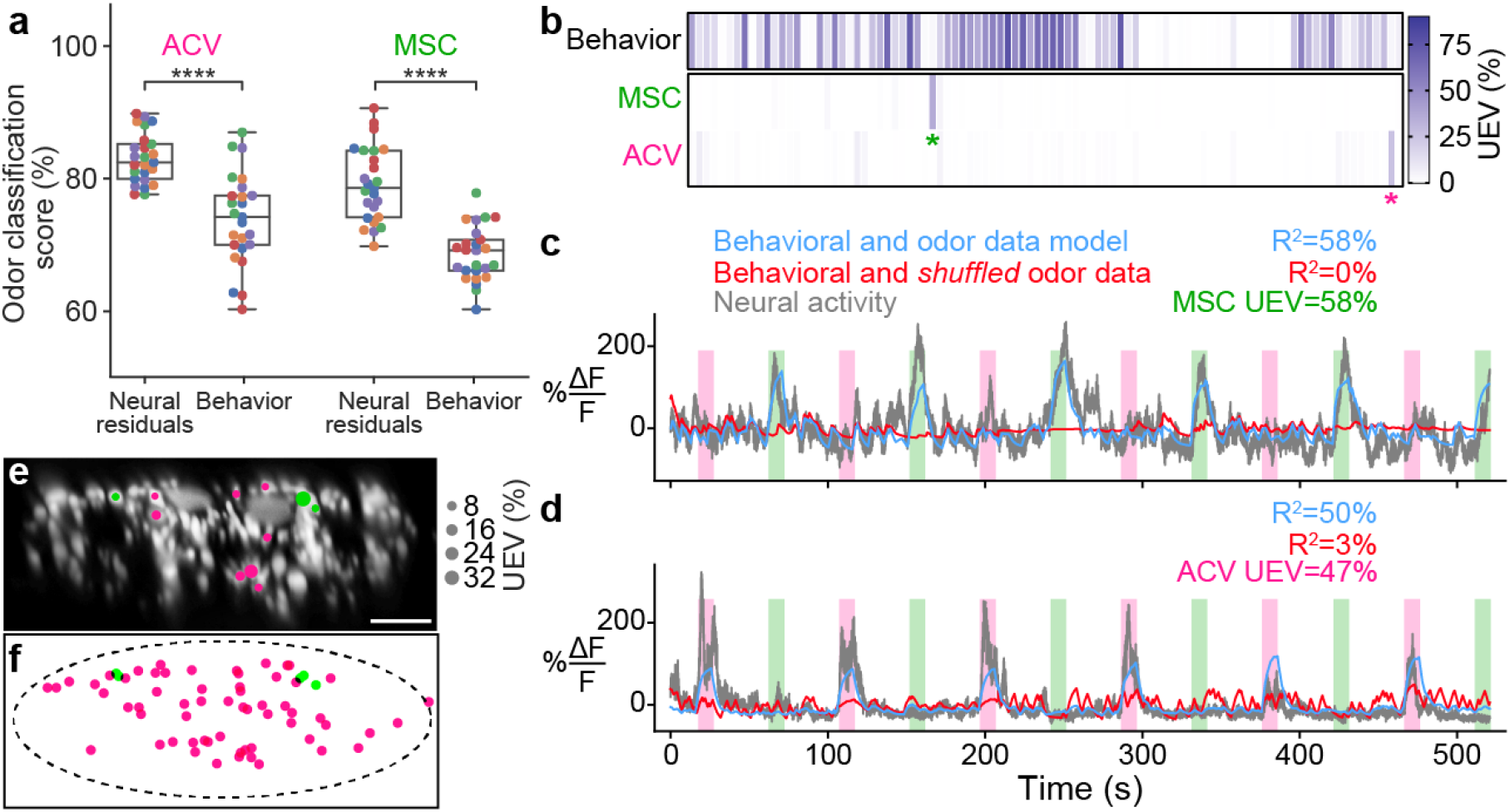
Odor encoding in descending neuron populations. **(a)** Neural residuals—obtained by subtracting convolved behavior regressors from raw neural data—can predict the presence of an odor significantly better than behavior regressors convolved with a calcium response function (‘Behavior’). Two-sided Mann-Whitney U test (**** indicates *p <* 10^−4^). The classification score was obtained using a linear discriminant classifier with cross-validation. Shown are five trials for five animals (colorcoded). **(b)** Matrix showing the cross-validated ridge regression unique explained variance (UEV) of a model that contains behavior and odor regressors for one animal. The first row (‘Behavior’) shows the composite *R*^2^ for all behavior regressors with odor regressors shuffled. The second and third rows show the UEVs for regressors of the odors methyl salicylate (MSC) or apple cider vinegar (ACV), respectively. Colored asterisks indicate neurons illustrated in panel **a. (c**,**d)** Example DNs best encoding **(c)** MSC or **(d)** ACV, respectively. Overlaid are traces of neural activity (gray), row one in the matrix (blue), and row one with odor data shuffled (red). **(e**,**f)** Locations of odor encoding neurons in **(e)** one individual, and **(f)** across all five animals. Scale bar is 10 µm.

### 2.8 Identifying individual descending neurons from population recordings

Until now, we have demonstrated that DN populations exhibit heterogeneous encoding: large, distributed groups encode walking and, by contrast, a few prominent pairs encode head grooming. Determining how these subpopulations control adaptive behavior is an important future challenge that will require a comprehensive approach examining phenomena ranging from global DN population dynamics down to the synaptic connectivity of individual DNs. The recent production of hundreds of sparse transgenic driver lines [10, 48] and connectomics datasets [22] suggest that this bridging of mechanistic scales may soon be within reach in *Drosophila*.

To illustrate how this might be accomplished, we aimed to identify specific DNs within our population imaging dataset. Specifically, while analyzing head grooming DNs, we noticed a large pair of ventral neurons **(Figure 5a)** that sometimes exhibited asymmetric activity **(Figure 5b, gray arrowheads)** when flies appeared to touch one rather than both antennae **(Video 5)**. To quantify this observation, we replayed limb 3D kinematics in NeuroMechFly, a biomechanical simulation of *Drosophila* [40] **(Figure 5c)**. By detecting leg-antennal collisions as a proxy for antenna deflection, we found that occasional asymmetries **(Figure 5d)** did coincide with asymmetric activity in corresponding neural data **(Figure 5b, purple traces)**. These results suggested that this pair of DNs encodes mechanosensory signals associated with antennal deflections.

**Figure 5:**
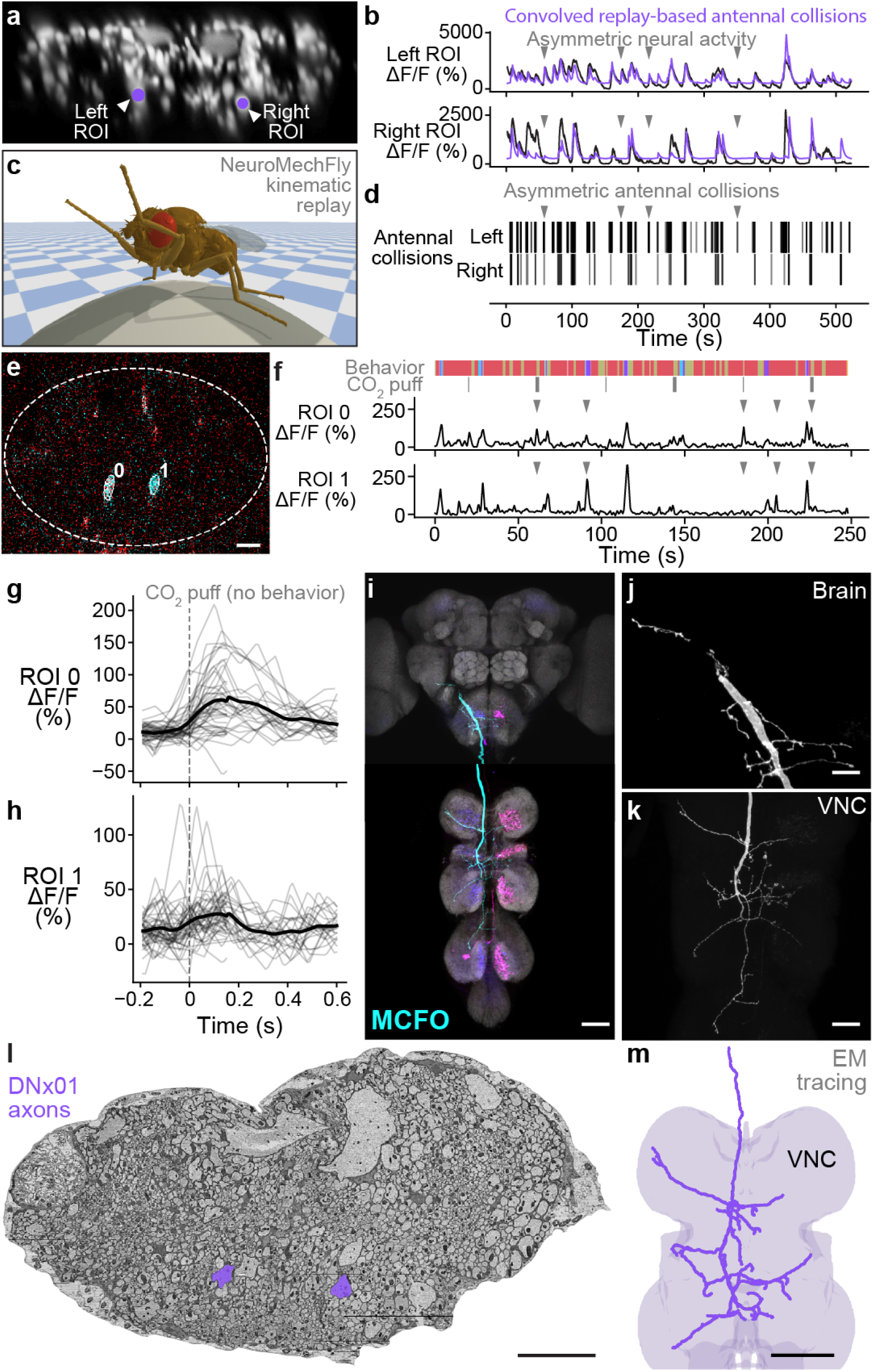
Identifying a pair of antennal deflection-encoding DNs from population recordings. **(a)** A pair of head groom-encoding DNs (purple circles and white arrowheads) can be identified from DN population recordings based on their shapes, locations, and activity patterns. **(b)** Example Δ*F/F* traces (black) of DNs highlighted in panel **a**. Sample time points with bilaterally asymmetric neural activity are indicated (gray arrowheads). Overlaid is a prediction of neural activity derived by convolving left and right antennal collisions measured through kinematic replay in the NeuroMechFly physics simulation (purple). **(c)** Kinematic replay of recorded joint angles in NeuroMechFly allow one to infer antennal collisions from real, recorded head grooming. **(d)** Left and right antennal collisions during simulated replay of head grooming shown in panel **b**. Sample time points with bilaterally asymmetric collisions are indicated (gray arrowheads). **(e)** Two-photon image of the cervical connective in a R65D11*>*OpGCaMP6f,tdTomato animal. Overlaid are ROIs identified using AxoID. The pair of axonal ROIs are in a similar ventral location and have a similarly large relative size like those seen in DN population recordings. Scale bar is 5 µm. **(f)** Sample neural activity traces from ROIs 0 and 1. Bilaterally asymmetric neural activity events (gray arrowheads), behaviors (color bar), and CO_2-_ puffs directed at the antennae (gray bars) are indicated. **(g**,**h)** CO_2_ puff-triggered average of neural activity for ROIs **(g)** 0 and **(h)** 1. Only events in which animals did not respond with head grooming or front leg rubbing were used. Stimuli were presented at *t* = 0. Shown are individual responses (gray lines) and their means (black lines). **(i)** Confocal volume z-projection of MCFO-expression in an R65D11-GAL4 animal. Cyan neuron morphology closely resembles DNx01 [10]. Scale bar is 50 µm. **(j**,**k)** Higher magnification MCFO image, isolating the putative DNx01 from panel **i**, of the **(j)** brain and **(k)** VNC. Scale bars are 20 µm. **(l)** The locations of axons in the cervical connective (purple) from neurons identified as DNx01. Scale bar is 10 µm. **(m)** Manual reconstruction of a DNx01 from panel **l**. Scale bar is 50 µm.

To further reveal the identity of these DNs, we examined data from our functional screen of sparse Gal4 and split Gal4 driver lines [49]. In this dataset, we observed similar asymmetric activity during antennal grooming in R65D11-GAL4. Coronal (x-z) two-photon imaging in R65D11 animals expressing OpGCaMP6f and tdTomato, shows axons that are similarly large and ventromedially located within the cervical connective **(Figure 5e)**. These also produce asymmetric activity during antennal grooming **(Figure 5f)**. This suggests that these neurons may report something unique to head grooming (e.g., coincident front limb movements) or simply antennal deflection. To distinguish between these possibilities, we analyzed neural responses to CO_2_ puff stimulation of the antennae, while discarding data with resulting head grooming or front leg rubbing to ensure that the antennae were not touched by the legs. We measured an increase in the activity of both DNs upon puff stimulation **(Figure 5g,h)** suggesting that, like the neurons recorded in DN populations, R65D11 neurons also encode sensory signals—antennal deflection—rather than behavior.

To confirm that these sparse neurons are DNs, we next performed MultiColor FlpOut (MCFO) [50] and confocal imaging of their morphologies. R65D11 drives expression in several neurons. However, we found similarly large axonal projections from only one set of neurons that descend from the brain to the VNC **(Figure 5i, cyan)**. Close examination of these neurites in the brain **(Figure 5j)** and VNC **(Figure 5k)** revealed a striking resemblance to the reported structure of DNx01 neurons [10] with cell bodies outside of the brain—putatively in the antennae and enabling antennal mechanosensing.

These results enable the analysis of synaptic connectivity in identified DNs. To illustrate this, based on their unique location and size, we identified DNx01s in a VNC electron microscopy dataset [22] **(Figure 5l)** via manual reconstruction and striking morphological similarity to R65D11 DNs **(Figure 5m)**. From this reconstruction, once the full VNC connectome becomes available, one may identify synaptic partners of DNx01 to further understand how they contribute to controlling antennal grooming and other behaviors. Taken together, these data suggest a possible road map for using functional, topological, and morphological data to decipher the cellular identity of individual DNs from population recordings.

## 3 Discussion

Here, by combining genetic and optical imaging approaches, we recorded the behavioral and sensory encoding of DN populations in behaving *Drosophila*. Across individual animals, we found that most recorded DNs encode walking. A smaller number are active during head grooming and resting. We did not find DNs encoding posterior movements, possibly due to the infrequency of this behavior. We also did not identify neurons that are active during multiple behaviors. This suggests that each ROI consists of individual neurons or that, if an ROI contains many neurons, they all show similar encoding. Subsets of walk-encoding DNs were also strongly active during turning: they were at higher density on the ipsilateral half of the cervical connective with respect to turn direction. By contrast, DNs distributed throughout the connective very weakly encoded walking speed. However, we caution that the small range of walking speeds in our data—flies accelerate rapidly from resting to walking and vice-versa— may mask stronger speed encoding. The partial overlap between turn- and speed-encoding DNs leaves open the possibility that neurons simultaneously encode these two properties in a differential steering fashion. Notably, we did not observe any DNs that are only active during transitions between multiple behaviors. However, the fly makes fast transitions. Thus, the signal-to-noise and temporal resolution of our approach may not be sufficient to identify such neurons—higher temporal resolution electrophysiological recordings would be required to confirm the absence of DN encoding for behavioral transitions or for precise limb kinematics.

The encoding—and presumptive control—of walking by large numbers of DNs support the notion that the brain tightly regulates locomotion. In contrast to walking, head grooming is encoded by fewer neurons in our data. This may be because grooming limb movements are more stereotyped and thus may only rely on controllers within the VNC (a notion that is supported by the ability of headless flies to perform spontaneous grooming [51]). Other studies have also shown brain-wide activity during walking but not during other behaviors [52–54]. This difference may arise because adaptive locomotion—to avoid obstacles [55], cross gaps [56], and court potential mates [57]—depends heavily on the brain’s descending signals. Thus, we hypothesize that, although a core set of command neurons can drive both walking and grooming, more DNs are additionally engaged during walking to allow for more flexible navigation in continuously changing and complex environments.

Because of the large number of DNs involved in walking, we hypothesized that subsets might represent parallel channels which are recruited depending on sensory context. For example, there may be DN subpopulations which drive walking in the presence of attractive odors and others engaged in the presence of aversive odors. This notion is supported by studies showing that optogenetic activation of a large variety of DNs elicits only a small set of stereotyped behaviors [17]. However, our population imaging data do not support this notion: largely the same DNs encode walking and head grooming irrespective of olfactory context.

A non-behavioral role for DNs has also been suggested by previous studies showing that the perception of a fictive odor modulates the activity of specific DNs involved in steering in immobile flies [29]. In line with this, we also identified DNs encoding odors and not behavior. Notably, the two odors we presented—apple cider vinegar and methyl salicylate—were encoded by distinct DNs. Extrapolating beyond ACV and MSC, it seems unlikely that DNs encode many individual odors with high specificity: the number of DNs is far smaller than what would be required to cover an enormous olfactory space. Instead, we speculate that DN odor-encoding may represent classes like attractive versus aversive odors or the valence of sensory inputs in general. Where odor information is conveyed within downstream motor circuits and for what purpose is a fascinating subject for future study.

Many fewer neurons encode head grooming as opposed to walking. Because our transgenic strain does not drive expression in SEZ neurons, we could not record several DNs whose activation has been shown to be sufficient to drive antennal grooming (aDN), front leg rubbing (DNg11), or both head grooming and front leg rubbing (DNg12) [20]. Thus, we expect that with the addition of these SEZ DNs the apparent dichotomy that many neurons encode walking and fewer encode grooming may become less pronounced. Nevertheless, groom-encoding DNs were notable in that they could be more reliably identified across individual animals. Among our head groom-encoding DNs, a pair passing through the ventral cervical connective appear to encode limb contact during antennal grooming as well as puff-dependent antennal deflections. We speculate that mechanosensory signals from the antennae may be used for feedback control in the VNC: being aware of whether the antennae are touched while grooming may allow for a continuous modulation of grooming kinematics and force application by the front legs. This may be a conserved control mechanism as similar DNs have been identified in the blow fly [58].

Our morphological and physiological evidence suggest that these antennal mechanosensory signals are provided by DNx01s, a subset of bilateral campaniform sensillum (bCS) neurons that are also found on the legs and form major presynaptic connections to fast motor neurons [22]. Thus, DNx01s may have a role beyond feedback control during grooming. This is also implied by the large size of DNx01 axons and their high sensitivity to puff-mediated antennal deflection. Because other DNs with large axons (e.g., giant fiber neurons) are often implicated in fast reflexive movements [59,60], DNx01s may be used to respond to, for example, strong gusts of wind that initiate a stance stabilization reflex.

Our work sets the stage for a more comprehensive, multi-scale investigation of how the brain regulates complex limb-dependent motor behaviors. Nevertheless, overcoming several technical limitations in our study should also be a focus of future work. First, although we could achieve precise limb kinematic measurements at 100 Hz, it will be critical to record neural data at equally high temporal resolution. The fly can walk with stride frequencies of up to 20 Hz [44] and leg movements during grooming occur at up to 7 Hz [43]. This currently makes it difficult to relate neural activity—read out by the relatively slow fluorescence fluctuations of a genetically-encoded calcium indicator—to individual joint angles and limb positions. To address this challenge, one might use faster indicators of neural activity (e.g., newer variants of GCaMP [61] or voltage indicators [62]). Additionally, coronal section two-photon imaging in the thoracic cervical connective with a piezo-driven objective lens limited our neural data acquisition to ∼16 Hz. One might perform data acquisition at a higher rate using more advanced imaging methods including single-objective light-sheet microscopy [63]. Alternatively, the fly could be forced to generate slower limb movements using a motorized treadmill [64]. Another challenge is that DNs from the subesophageal zone (SEZ) are absent in our driver line. The SEZ is considered a center for action section [65, 66] and is known to house numerous DNs [10, 11]. Thus, complementing our driver line with an SEZ-expressing transgene [67] would enable a fully comprehensive recording of DN population dynamics. Nevertheless, we expect our observation to remain intact: locomotion is regulated by large DN populations in a distributed manner, while more stereotyped grooming behaviors engage fewer DNs. This would suggest a dichotomy in DN population control for flexible versus stereotyped motor behaviors. Future studies may test if this holds true as well for wing-dependent behaviors like continuous steering during flight [13, 16] versus stereotyped wing displays during courtship [68].

## 5 Materials and Methods

### 5.1 Fly husbandry and stocks

All flies were kept at 25 °C and 50 % humidity on a 12 h light-dark-cycle. Flies were 10 days posteclosion (dpe) for experiments, and had been starved overnight on a wet precision wipe (Kimtech Science, 05511, USA). Sources of each genotype used are indicated in **Table 1**.

### 5.2 Olfactometer

Mass flow controllers (MFC) were used to regulate air flow (Vögtlin, GSC-B4SA-BB23 (2 L min^−1^), GSC-A3KA-BB22 (100 mL min^−1^)). The larger MFC, set to 42 mL min^−1^, was used to continuously bubble odor vials, maintaining a constant head space odorant concentration. The smaller MFC was used to stimulate the fly at 41 mL min^−1^. We directed air flow using six solenoid valves (SMC, S070C-6AG-32, Japan) controlled by an Arduino Mega (Arduino, A000067, Italy). One valve in front of each of the three odor vials was used to switch between inputs from each MFC. A second valve after each odor vial was used to direct air flow either towards the fly or into an exhaust. The second valve was placed as close to the fly as possible (∼10 cm) to minimize the delay between solenoid switching and odor stimulation. To direct air flow to the fly’s antennae, we used a glass capillary held in place by a Sensapex zero-drift micro-manipulator (Sensapex, uMp-3, Finland). We minimized mechanical perturbations by blowing humidified (non-odorized) air onto the fly between odor trials. We used a PID (Aurora Scientific, miniPID, Canada) and visual assessment of animal behavior to confirm that switching generated minimal mechanical perturbations.

### 5.3 Two-photon microscopy

We performed neural recordings using a ThorLabs Bergamo two-photon microscope (Thorlabs, Bergamo II, USA) connected to a Mai Tai DeepSee laser (Spectra Physics, Mai Tai DeepSee, USA). To perform coronal section imaging, we operated the microscope in kymograph mode using the galvo-resonant light path. Images were acquired at a magnification of 7.4 X, resulting in a 82.3 µm wide field-of-view (FOV). To prevent the cervical connective from leaving the FOV, the full range of a 100 µm piezo collar was used to scan the objective lens (Olympus XLUMPlanFLN 20 X, 1.0 NA with 2 mm working distance) in the z-axis. We could achieve a frame rate of approximately 16 Hz by sampling 736 × 480 pixel (x-and z-axes, respectively) images and by enabling bidirectional scanning, with only one fly-back frame.

### 5.4 Neural recordings

#### 5.4.1 Descending neuron population recordings

Flies were dissected to obtain optical access to the thoracic cervical connective, as described in [14]. Briefly, we opened the cuticle using a syringe and waited for the flight muscles to degrade before resecting trachea, the proventriculus, and the salivary glands. After removing these tissues covering the VNC, an implant [31] was inserted into the thoracic cavity to prevent inflation of the trachea and to ensure a clear view of the cervical connective for extended periods of time. Flies were then given several minutes to adapt to positioning over a spherical treadmill in the two-photon microscope system. During this adaptation period, the nozzle of the olfactometer was positioned approximately 2 mm in front of the the fly’s head. As well, the thoracic cervical connective was brought into the imaging FOV. Following alignment, their position was further adjusted to maximize fluorescence signal and to minimize the possibility of neurites leaving the imaging FOV. For each fly, a minimum of five 9 min trials were recorded using a laser power of 9.25 mW at 930 nm.

#### 5.4.2 Sparse DNx01 recordings

Recordings of DNx01s in the R65D11-GAL4 driver line were performed as described in [49]. This differed only slightly from DN population recordings in the following ways. First, flies were not starved. Second, the Gal4 drove expression of OpGCaMP6f rather than GCaMP6s. Third, a thoracic implant was not used. Fourth, instead of being presented with a constant flow of air and odors, animals were stimulated with CO_2_ puffs of alternating length (0.5 s, 2 s) spaced 40 s apart. A higher pixel dwell time was used to achieve acceptably high imaging signal-to-noise. This resulted in a slower two-photon image acquisition rate (4.3 fps) which was matched by a slower behavior recording frequency (30 Hz). For additional details and a description of the stimulation system see [49]. Note that images of the connective in **Figure 5a** and **Figure 5e** appear to have different heights due to a difference in z-step size during image acquisition.

### 5.5 Post-processing of two-photon imaging data

#### 5.5.1 Descending neuron population recordings

Binary output files from ThorImage (Thorlabs, ThorImage 3.2, USA) were converted into separate tiff files for each channel using custom Python code (https://doi.org/10.5281/zenodo.5501119). Images acquired from the red channel were then denoised [69] and two-way alignment offset was corrected (https://doi.org/10.5281/zenodo.6475468) based on denoised images. We then used optic flow estimation to correct image translations and deformations based on the denoised red channel images (https://doi.org/10.5281/zenodo.6475525). The green channel was then corrected based on the motion field estimated from the red channel. Finally, we trained a DeepInterpolation network [37] for each fly using the first 500 motion corrected green channel images from each experimental trial (batch size=20; epochs=20; pre-post frames=30). The first and the last trials were used as validation datasets. The trained network was then used to denoise green channel images.

Baseline fluorescence was then determined on a fly-wise and pixel-wise basis across all trials. The baseline of a pixel was defined as the minimum ‘mean of 15 consecutive values’ across all experimental trials. Motion correction introduces zeroes to two-photon images in background regions that were out of the FOV prior to warping. Therefore, values close to zero (floating point inaccuracy) were set to the maximum of the datatype of the array. This means essentially ignoring these pixels and their immediate surroundings for baseline computations. We used the baseline image *F*_0_ to calculate 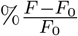 images. These images were only used for visualization. To identify ROIs/neurons, we created a maximum intensity projection of 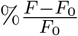 images and manually annotated ROIs. The 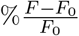 of each ROI was computed by first spatially averaging it’s raw pixel values. We then calculated the baseline of this average as described for a single pixel above. For brevity and readability, we refer to 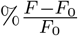 as %Δ*F/F* throughout the manuscript.

#### 5.5.2 Sparse DNx01 descending neuron recordings

ROIs were detected using AxoID. Raw, non-denoised traces were used for analysis. For more details concerning data post processing see [49].

### 5.6 Behavior classification and quantification

#### 5.6.1 Behavior measurement system

The behavior of tethered animals was recorded using a previously described [33] 7-camera (Basler, acA1920-150um, Germany) system. Animals were illuminated using an infrared (850 nm) ring light (CSS, LDR2-74IR2-850-LA, Japan). To track the joint positions of each leg, six cameras were equipped with 94 mm focal length 1.00 X InfiniStix lenses (Infinity, 94mm/1.00x, USA). All cameras recorded data at 100 fps and were synchronized using a hardware trigger. For more details see [33].

#### 5.6.2 Inferring fictive locomotor trajectories

Video data from the front camera were processed using FicTrac [34] to track ball movements. This camera was outfitted with a lens allowing adjustable focus and zoom (Computar, MLM3X-MP, 0.3X- 1X, 1:4.5, Japan). To improve tracking accuracy, the quality factor of FicTrac was set to 40. The circumference of the ball was detected automatically using a Hough circle transform on the mean projection of all images for a given experimental trial. To determine the vertical angular FOV, *α*, the value given in the specifications of the Computar lens (8.74°) had to be adjusted, accommodating a smaller sensor size. We first determined the focal length to be 43.18 mm using Equation 1, where *H* is the height of the sensor. This was set to 6.6 mm for a 2/3” sensor.

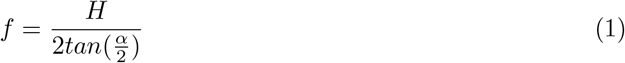

The ROI of the Basler camera was set to 960×480 pixels, reducing the effective sensor height from 5.8 mm to 2.32 mm. Rearranging Equation 1 and plugging in *f* and *H* yields a vertical angular FOV of 3.05°. Since the camera was already aligned with the animal, the camera-to-animal transform was set to zero. To obtain the fly’s trajectory, we developed custom code that integrates rotational velocities (https://github.com/NeLy-EPFL/utils_ballrot).

#### 5.6.3 Postprocessing of 3D pose estimates

Outliers in 3D pose data were corrected as described in [49]. Briefly, we detected outliers based on changes in leg segment length and chose the pair of cameras with minimal reprojection error for triangulation. After outlier correction, the data were aligned and joint angles were computed using published code [40]: https://github.com/NeLy-EPFL/df3dPostProcessing/tree/outlier_correction.

#### 5.6.4 Classification of behaviors

Behaviors were classified based on limb joint angles using the approach described in [41]. Briefly, a network was trained using 1 min of annotations for each fly and heuristic labels. Motion energy, ball rotations, and joint positions where used to generate the heuristic labels. To compute the motion energy, each joint position was convolved with the finite difference coefficients of length nine, estimating the first derivative. After computing the *L*^1^-norm, the signal was filtered with a 10th order low-pass Butterworth filter of critical frequency 4 Hz. The total, front, and hind motion energy were computed by summing over all joints, all front leg joints, and all hindleg joints, respectively. Forward ball rotation speeds were processed using the same Butterworth filter described above. First, we assigned heuristic labels for walking by thresholding the filtered forward walking velocity at 0.5 mm s^−1^. The remaining frames with a motion energy smaller than 0.3 were then classified as resting. Next, heuristic labels for front movements (front motion energy *>* 0.2 and hind motion energy *<* 0.2) and posterior movements (front motion energy *<* 0.2 and hind motion energy *>* 0.2) were assigned to all remaining frames. The front movement labels were further split into head grooming and front leg rubbing by thresholding the height of the front leg tarsi (the average between left and the right tarsi) at 0.05. After each step, a hysteresis filter was applied. This filter only changes state when at least 50 consecutive frames are in a new state. Based on a hyperparameter search using ‘leave one fly out’ cross validation on the hand labels **(Figure S1g)**, we selected the weights of λ_*pred*_ = 0 and λ_*weak*_ = 1 for the loss.

#### 5.6.5 Biomechanical simulation and antennal collision detection

To infer limb-antennal collisions, we performed kinematic replay using the NeuroMechFly physics simulation framework as described in [40]. We used joint angles to replay real limb movements in the simulation. To avoid model explosion and accumulating errors over long simulation times, we ran kinematic replay on time segments shorter than the full experimental trials. These segments consisted of individual head grooming and front leg rubbing events. The default head angle was fixed to 20°. The aristae yaw values were set to −32° and 32° and pedicel yaw values were set to 3−3° and 33° for the left and right sides, respectively. To speed up the simulation, we only detected collisions between the front legs and head segments.

### 5.7 Confocal imaging of the brain and ventral nerve cord

Confocal images were acquired using a Zeiss LSM700 microscope. These images were then registered to a brain and VNC template described in [49] using the method from [70]. Brain and VNC sample preparation was performed as described in [49]. Both primary and secondary antibodies were applied for 24 h and the sample was rinsed 2-3 times after each step. Antibodies and concentrations used for staining are indicated in **Table 2**.

### 5.8 Electron microscopy identification and tracing

Within an electron microscopy dataset of the ventral nerve cord and neck connective [22], we identified the pair of DNx01s based on their large-caliber axons in the cervical connective positioned ventral to the giant fiber neurons axons (**Figure 5l**). We then manually reconstructed all branches of one DNx01 neuron using CATMAID [71, 72] as described in [22]. The reconstructed neuron was then registered to the female ventral nerve cord standard template [73] using an elastix-based atlas registration as described in [22].

### 5.9 Data analysis

#### 5.9.1 Linear regression modeling

We relied primarily on regression techniques to link behavioral and neural data. Here we first describe the general approaches used and then discuss details and modifications for individual figure panels. To evaluate the success of regression models, we calculated explained variance in the form of the coefficient of determination (*R*^2^) and unique explained variance (UEV) [74]. The explained variance is a measure of how much additional variance is explained by the model compared to an intercept only model (i.e., approximating the data by taking the mean). A definition of the coefficient of determination can be found in Equation 2, where SSE is the sum of squares of the error, SST is the total sum of squares, *y*_*i*_ is a data point, *ŷ*_*i*_ is the prediction of *y*_*i*_, and 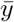 is the mean of all data points.

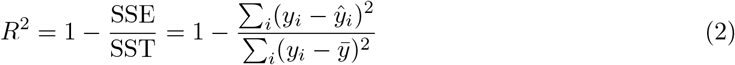

Note that, *R*^2^ becomes negative when SSE is larger than SST. This is the case when the model prediction introduces additional variance due to overfitting. UEV is a measure for the importance of individual or a subset of regressors. It is computed as the reduction in *R*^2^ when a particular subset of regressors is randomly shuffled (Equation 3).

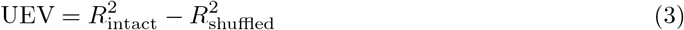

We performed 5-fold cross-validation for all of our regression results to ensure good generalization. Non-regularized linear regression sometimes led to overfitting and negative *R*^2^ values. Therefore, we used ridge regression. The ridge coefficient was determined using 5-fold nested cross-validation on the training data set. In some cases we still observed small negative *R*^2^ values after applying ridge regularization. These were set to zero, in particular to avoid problems when computing UEVs. To account for the long decay dynamics of our calcium indicator we convolved behavior variables with an approximation of the calcium response function (crf) (Equation 4).

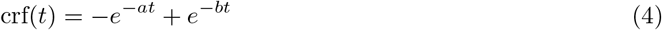

We used *a* = 7.4 and *b* = 0.3 to approximate the rise and decay times, respectively, as reported in [35]. We also normalized the function to integrate to one on the interval 0–30 s.

##### Figure 2 & Figure S3

In these figures we predicted behavior from neural activity. To accomplish this, we trained models for all pairs of behaviors and neurons (e.g., walking and ROI 41 for the upper plot of **Figure 2a**). The target variable is a binary variable indicating whether the fly is walking or not. This was convolved with the crf (black line in panel **a**). The single regressor besides the intercept in the model is the Δ*F/F* of a single neuron. **Figure 2b** and **Figure S3c** show the *R*^2^ values for all models. Each neuron was then assigned to be encoding the behavior with the maximum *R*^2^. For neurons with maxima smaller than 5%, no behavior was assigned.

##### Figure S2

Using the approach described in the previous section, we observed that some neurons appear to predict both walking and posterior movements, while others predict both head grooming and front leg rubbing. Because fluorescence decays slowly following the cessation of neural activity, this may be caused by the frequent sequential occurrence of two behaviors. For instance, if front leg rubbing often occurs after head grooming, the Δ*F/F* of a head groom encoding neuron may still be elevated during front leg rubbing. This can lead to false positives in our analysis. To address this potential artifact, we predicted neural activity from multiple behavior regressors (i.e., binary behavior indicators convolved with the crf). We then calculated the UEV for each behavior regressor. For example, when the front leg rubbing regressor is shuffled the *R*^2^ will not decrease by much because the head grooming regressor includes the expected decay through convolution with a crf. The model of **Figure S2a** includes a walking and a posterior movements regressor. For **Figure S2b** the model includes a head grooming and a front leg rubbing regressor. No other regressors were included in these models and for a given model, only data during one of these two behaviors were used.

##### Figure 3 & Figure S7

For **Figure 3a** we used all behavior regressors and the Δ*F/F* of all neurons to predict ball rotation speeds convolved with a crf. We then temporally shuffled each of the neural regressors to calculate their UEVs. Since knowing whether the fly is walking or not provides some information about forward speed, the 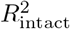 did not decrease to zero in the speed prediction (panel **a**, 2nd row). We address this issue in panel **b** by only including walking frames but otherwise using the same approach as in panel **a**. To pinpoint the encoding of ball rotations to individual neurons **(Figure 3c)**, we made two changes to our approach. First, instead of predicting turning in general, we split the turning velocity into right and left turning. Second, we only included the Δ*F/F* of a single neuron in the model rather than the data from all neurons. As before, the target variables were spherical treadmill rotation speeds—right turning, left turning, and forward walking—convolved with the crf.

##### Figure 4 & Figure S8

To model odor encoding, we predicted the activity of a single neuron using behavior regressors, including crf-convolved spherical treadmill rotation speeds, and odor regressors constructed by convolving the crf with a binary variable indicating whether a given odor was present or not. To determine how much neural variance can be explained by the behavior regressors in our model, we shuffled both odor regressors **(Figure 4b, top row)**. We then calculated the UEV for each odor by first computing the *R*^2^ of the model with all regressors intact, and then subtracting the *R*^2^ of the model after shuffling the odor regressor in question **(Figure 4b bottom)**. In panels **c** and **d**, the intact model’s prediction (blue) and the shuffled model’s prediction (red) are shown.

##### Figure S9

To see if a neuron’s behavior encoding could change depending on the context of which odor is present, we only examined the most commonly encoded behaviors: walking and head grooming. Each row is equivalent to the the walking or head grooming rows in **Figure S3b**. However, here we only used subsets of the data to train and validate our models. Each row only uses data when one odor (or humidified air) is present. If there is no context dependence, each of the three rows should look identical. However, we note that due to the significant reduction in the amount of data for each model, noise can introduce variation across conditions. Humidified air data was subsampled to match the amount of data available for the odors MSC and ACV. To test whether the encoding was significantly different across contexts, we used a two-sided Mann-Whitney U test. The data points are individual cross-validation folds from each trial. Elsewhere in this study we report cross-validation means.

##### Figure 5

In panel **b** we perform a regression to predict one neuron’s activity using the intercept as well as a crf-convolved antennal collision regressor derived from data in panel **d**.

#### 5.9.2 Kernel density estimation

We performed 2D kernel density estimation **(Figure 2h,i)** using SciPy’s gaussian kde [75]. We performed 1D kernel density estimation **(Figure 3h,j; Figure S7d,f)** using sklearn [76]. We normalized kernel density estimates to correct for the variable density of neurons across the connective. The normalization factor was computed as a kernel density estimate of all annotated neuron locations. The kernel bandwidth was determined using leave-one-out cross-validation. This maximizes the log-likelihood of each sample under the model. Data from all flies were used to determine the bandwidth.

#### 5.9.3 Principal component analysis

We performed principal component analysis (PCA) on DN population data. First, as in [45], we observed that PCs from time derivatives of fluorescence traces produce more organized state space trajectories. Therefore, we calculated the derivative of the Δ*F/F* traces for each neuron (extracted from non-denoised imaging data) using total variation regularization [77]. Empirically, we found that a regularization parameter of 5000 strikes a good balance between bias and variance in the derivatives. We then performed PCA on the derivatives of all neural data during walking only. This allowed us to specifically ask whether, during walking, neural activity largely remained constant or diverged as a function of specific locomotor features. We then embedded walking and resting neural data epochs into the same PC space (i.e., we did not refit the PCs for resting). We visualized the loadings of individual ROIs/neurons using vectors to illustrate how these neurons influence the position in PC space. For clarity we only show vectors for whom 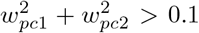, where *w*_*pc*1_ and *w*_*pc*2_ are the loadings of the first and second principal components.

#### 5.9.4 Linear discriminant analysis

We trained linear discriminant models to distinguish between ACV and humidified air, as well as MSC and humidified air **(Figure 4a)**. To evaluate classification accuracy, the classification score was cross-validated. Input data to the model were either behavior regressors or neural residuals. The residuals were computed using ridge regression (as described above) with behavior regressors as input. In both cases the behavior regressors were convolved with the crf.

#### 5.9.5 Event-triggered averaging

Here, ‘events’ describe individual epochs of a particular behavior. To perform event-triggered averaging of images/videos **(Video 2-4)**, we first identified all events that had no similar event in the previous 1 s. Raw microscope recordings were then chopped into blocks starting 1 s prior to event onset and lasting 4 s after event onset or until the end of the event, whichever was shorter. Each block was then converted into Δ*F/F* using the mean of the first five frames as a baseline. All blocks were then temporally aligned and averaged frame-by-frame. We discarded behavior videos with fewer than 50 events 1 s after event onset. Event-triggered averaging of neural traces **(Figure 5g,h)** was performed in a similar fashion. However, instead of using raw images, Δ*F/F* traces were used and no block-wise Δ*F/F* was computed. The values were then averaged one time-point at a time.

#### 5.9.6 Correlation coefficients

We calculated the Pearson’s correlation coefficient for each trial between the raw Δ*F/F* trace and a time shifted Δ*F/F* trace calculated using denoised images **(Figure S1b)**. We then either grouped the values by time lag and averaged each group (bottom), or we found the time shift with maximal cross-correlation. In **Figure 3d** and **Figure S7b** we perform several types of correlation analyses. First, the larger matrices show the correlation between neurons. This is computed as the Pearson’s correlation coefficient between the Δ*F/F* values of a pair of neurons. Correlations between neural activity and turning or walking speed were calculated as the Pearson’s correlation coefficient between the Δ*F/F* values of a neuron and corresponding spherical treadmill rotations, using only data when the fly was classified as walking. All of the above calculations were performed on individual trials and then averaged across all trials.

#### 5.9.7 Hierarchical clustering

We used Ward’s method [78] to hierarchicaly cluster and sort neurons based on their correlation **(Figure 3d)**. The distance between pairs of neurons was set to 1 − *r*, where *r* is the Pearson’s correlation coefficient for the two neurons.

## 6 Supplementary Tables

**Table 1:** Sources of transgenic *Drosophila melanogaster* strains.

| Genotype | Source | Reference |
| --- | --- | --- |
| w; $\frac{\text{tub>stop>GAL80}}{(\text{CyO})}$; $\frac{\text{MKRS}}{\text{TM3, Sb (21-21)}}$ | Asahina lab<br>(Salk Institute,<br>San Diego, CA USA) | [32] |
| w; $\frac{\text{Otd-nls:FLP (attP40)}}{(\text{CyO})}$; $\frac{\text{TM2}}{\text{TM6B(26-27)}}$ | Asahina lab<br>(Salk Institute,<br>San Diego, CA USA) | [32] |
| w; tubP-(FRT.GAL80); | Bloomington Drosophila<br>Stock Center (#62103) |  |
| w; + ; R57C10-GAL4 | Bloomington Drosophila<br>Stock Center (#39171) | [48] |
| w; + ; 10xUAS-IVS-myr::smGFP-FLAG (attP2) | Bloomington Drosophila<br>Stock Center (#62147) | [50] |
| + [HCS]; P20XUAS-IVS-GCaMP6s attP40;<br>Pw[+mC]=UAS-tdTom.S3 | Dickinson lab<br>(Caltech,<br>Pasadena, CA USA) |  |
| ;P20XUAS-IVS-Syn21-OpGCaMP6f-464 p10 su(Hw)attP5;<br>Pw[+mC]=UAS-tdTom.S3 | Dickinson laboratory<br>(Caltech,<br>Pasadena, CA USA) |  |
| R57C10-446 Flp2::PEST in su(Hw)attP8; ; HA-V5-FLAG | Bloomington Drosophila<br>Stock Center (#64089) | [50] |
| w;;P{w[+mC]=20xUAS-DSCP><br>H2A::sfGFP-T2A-mKOk::Caax} JK66B | McCabe lab<br>(EPFL,<br>Lausanne, Switzerland) | [79] |

**Table 2:** Antibodies and concentrations used to stain *Drosophila melanogaster* nervous tissues.

| Produced in | Antibody | Dilution | Manufacturer | RRID |
| --- | --- | --- | --- | --- |
| rabbit | anti-GFP | 1:500 | Thermofisher | AB 2536526 |
| mouse | anti-Bruchpilot | 1:20 | Developmental Studies<br>Hybridoma Bank | AB 2314866 |
| goat | anti-rabbit<br>conjugated with Alexa 488 | 1:500 | Thermofisher | AB 143165 |
| goat | anti-mouse<br>conjugated with Alexa 633 | 1:500 | Thermofisher | AB 2535719 |
| rabbit | anti-HA-tag | 1:300 | Cell Signaling Technology | AB 1549585 |
| rat | anti-FLAG-tag DYKDDDDK | 1:150 | Novus | AB 1625981 |

## 7 Supplementary Figures

**Figure S1:**
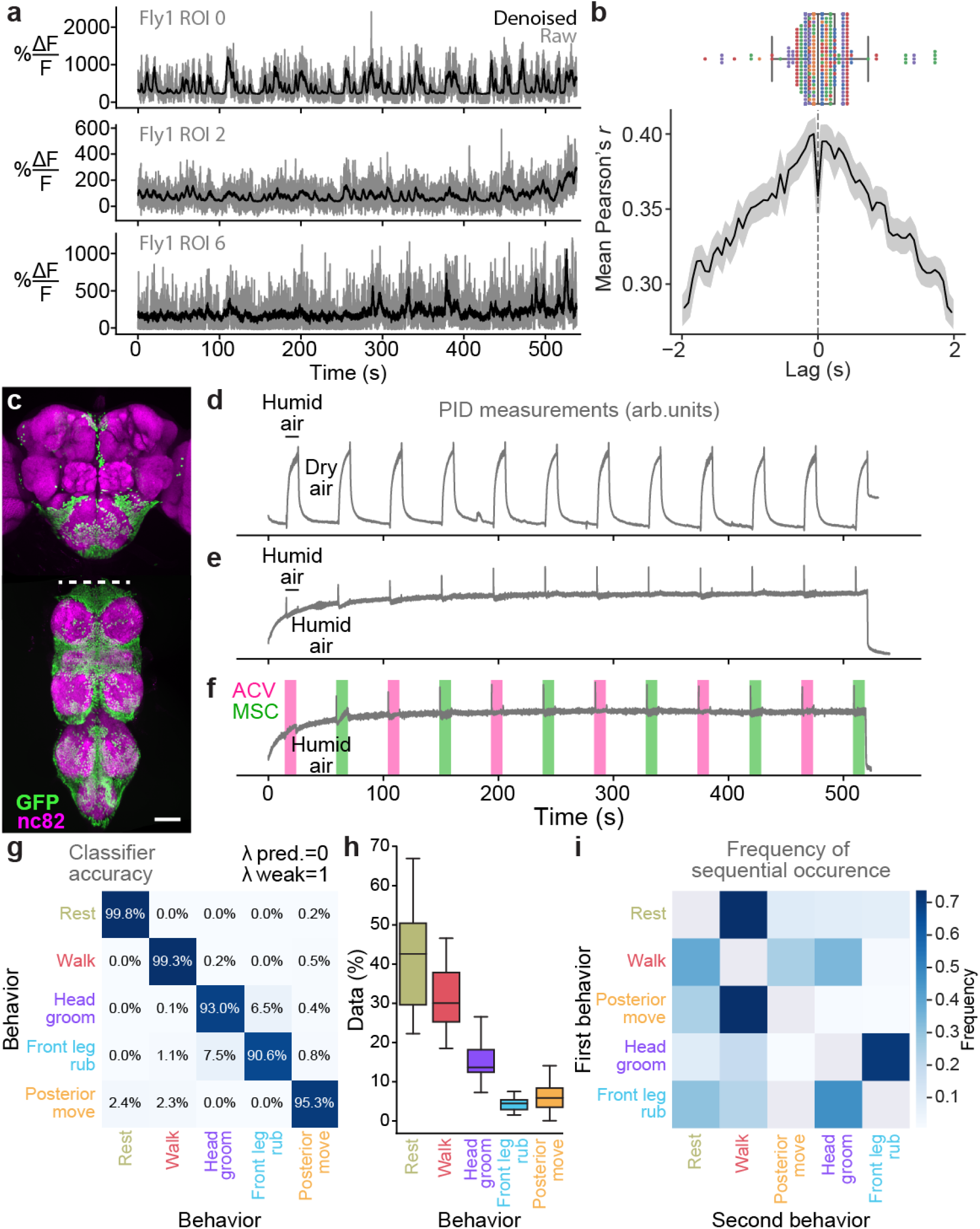
Supporting details for neural denoising, driver line expression, odor stimulation, and behavior quantification. **(a)** Example Δ*F/F* traces extracted from optic-flow registered ‘Raw’ (gray) and corresponding ‘Denoised’ (black) images. **(b, top)** Lag between denoised and raw Δ*F/F* traces with maximal cross-correlation. Overlaid are individual data points from each fly (colorcoded). **(b, bottom)** Average cross-correlation for all ROIs and flies (solid line) and corresponding 95% confidence interval (shaded region). **(c)** Z-projected confocal image of the genetic complement of our ‘brain only’ driver line (otd-nls:FLPo,tub*>*stop*>*GAL80; R57C10-GAL4) expressing nuclear GFP. Shown are neuropil (‘nc82’, magenta) and GFP (green) staining. Scale bar is 50 µm. **(d-f)** Photoionization detector (PID) measurements during our odor delivery protocol switching between **(d)** humid and dry air, **(e)** humid and humid air (to measure valve-related transients) or **(f)** humid air, MSC, and ACV odors. **(g)** Results of leave-one-fly-out cross-validation hyperparameter search for our behavior classifier. The values λ_*weak*_ = 1 and λ_*pred*_ = 0 yield the highest classification accuracy. **(h)** Relative frequency of classified behaviors in our dataset (*n*=5 animals). **(i)** The frequency of transitions between sequential behaviors.

**Figure S2:**
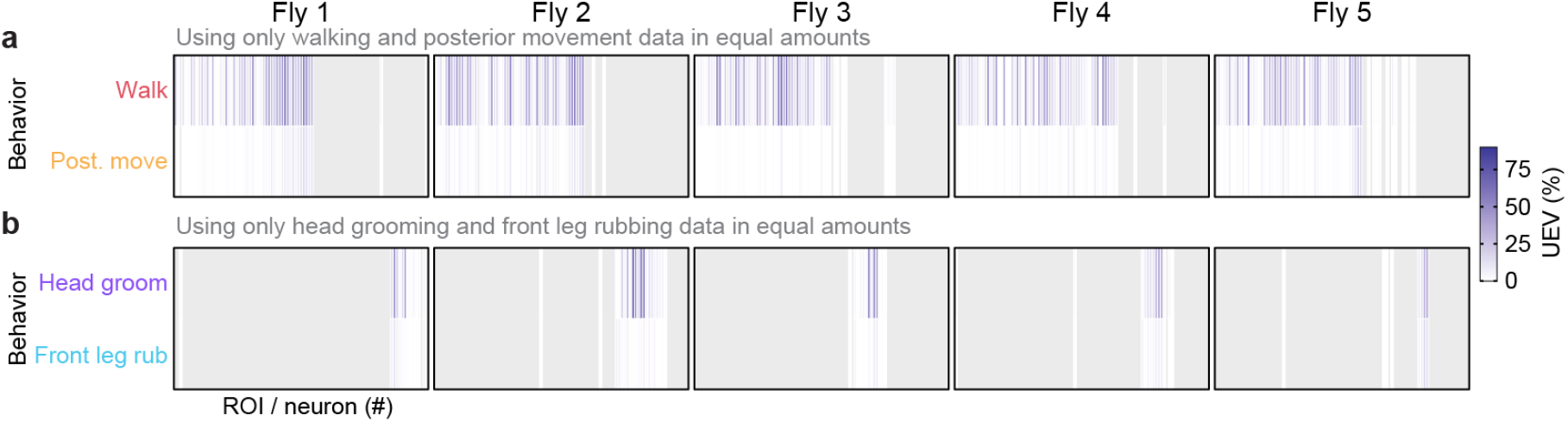
Disentangling the relative encoding of frequently sequential behavior pairs. **(a-b)** Cross-validation mean of neural variance uniquely explained by **(a)** walking versus posterior movements or **(b)** head grooming versus front leg rubbing. In both cases only data acquired during the two compared behaviors were analyzed. Additionally, data were balanced to have an equal amount across both behaviors.

**Figure S3:**
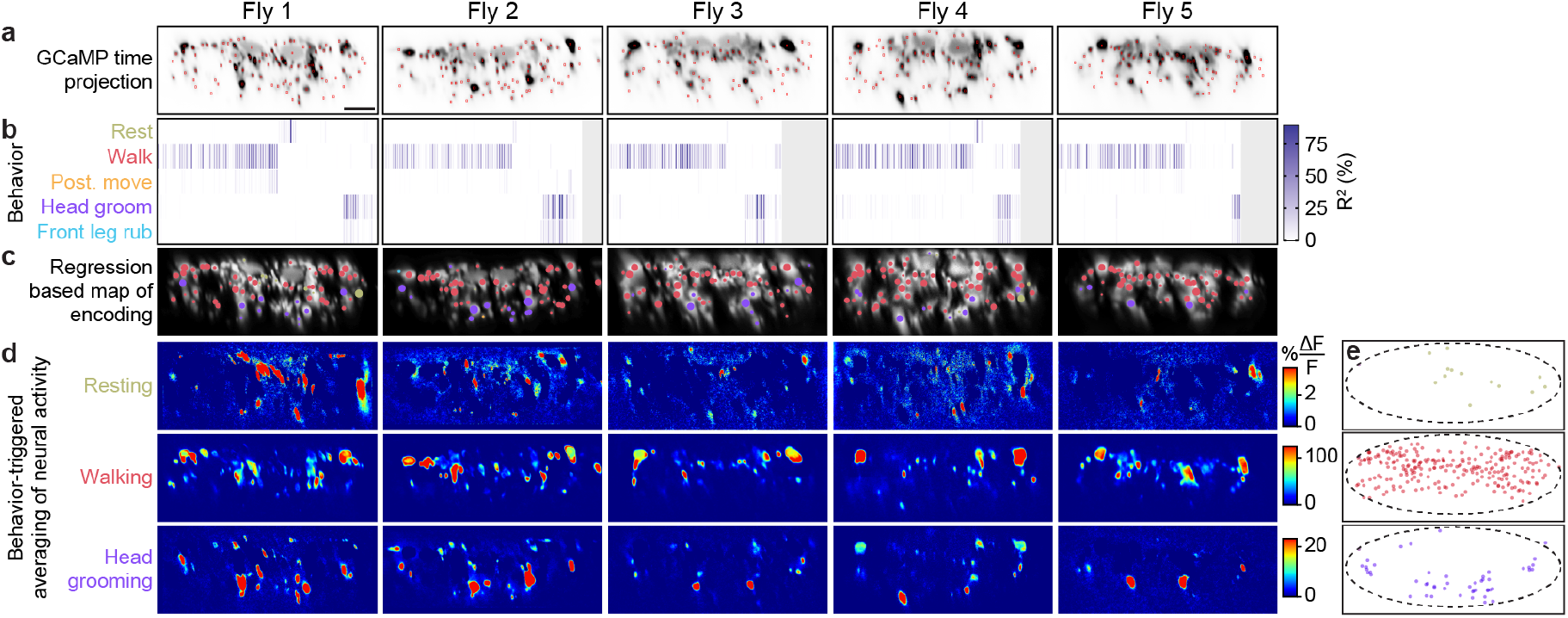
Encoding of behavior in descending neuron populations across individual animals. **(a)** Mean time projections of GCaMP6s fluorescence over a nine minute recording for five animals. Images are inverted for clarity, illustrating high mean fluorescence (black). Manually identified DN regions-of-interest (ROIs) are shown (red rectangles). Scale bar is 10 µm. All subpanels and panels **c-d** share the same scale. **(b)** The cross-validation mean of behavioral variance explained by DNs for each animal (Fly 1: *n*=95 ROIs; Fly 2: *n*=86 ROIs; Fly 3: *n*=75 ROIs; Fly 4: *n*=81 ROIs; Fly 5: *n*=79 ROIs). **(c)** DNs color-coded by the behavior their activities best explain. Radius scales with the amount of variance explained. **(d)** Behavior-triggered average Δ*F/F* images for the most common behaviors—resting, walking, and head grooming. **(e)** Locations of DNs for the classified behavior they encode best. Data are pooled across five animals.

**Figure S4:**
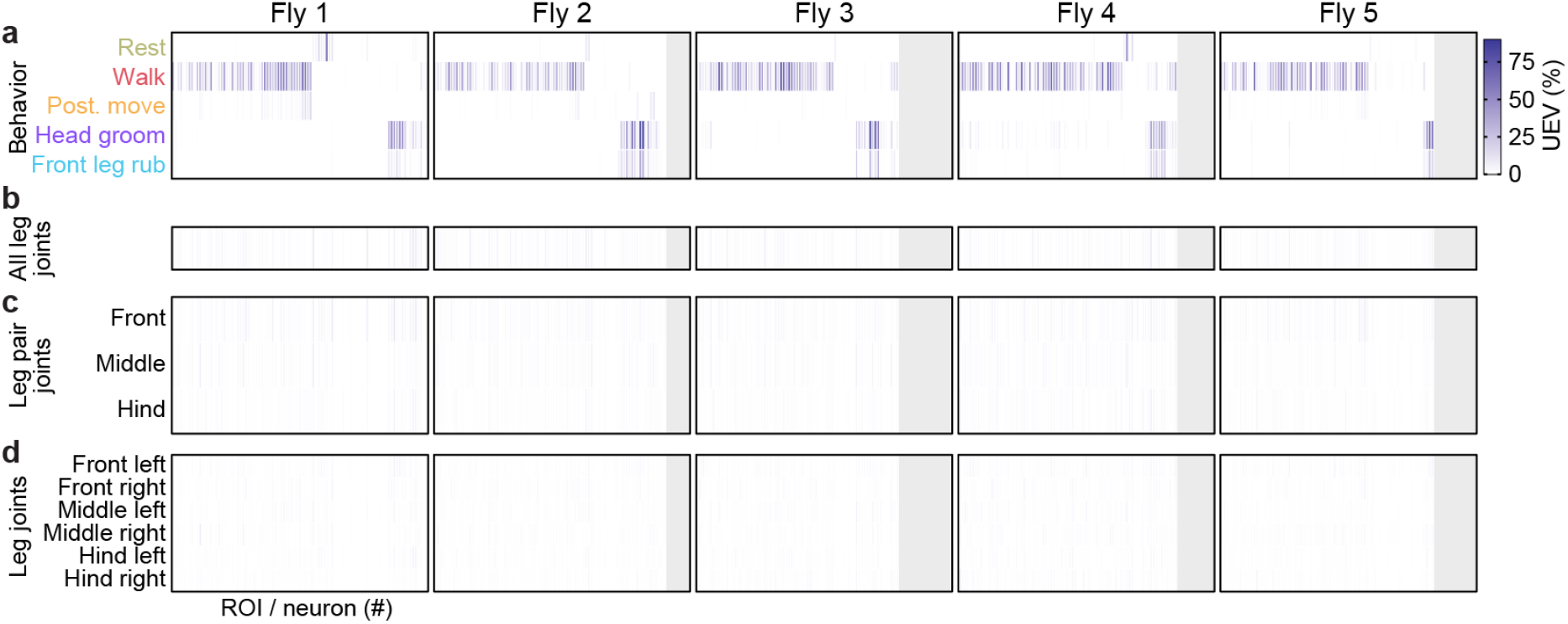
Neural variance explained by distinct kinematic features. **(a-d)** Amount of neural variance that can be uniquely explained by **(a)** classified behaviors (taken from the previous figure) **(b)** all joint movements **(c)**, leg pair movements, or **(d)** individual leg movements.

**Figure S5:**
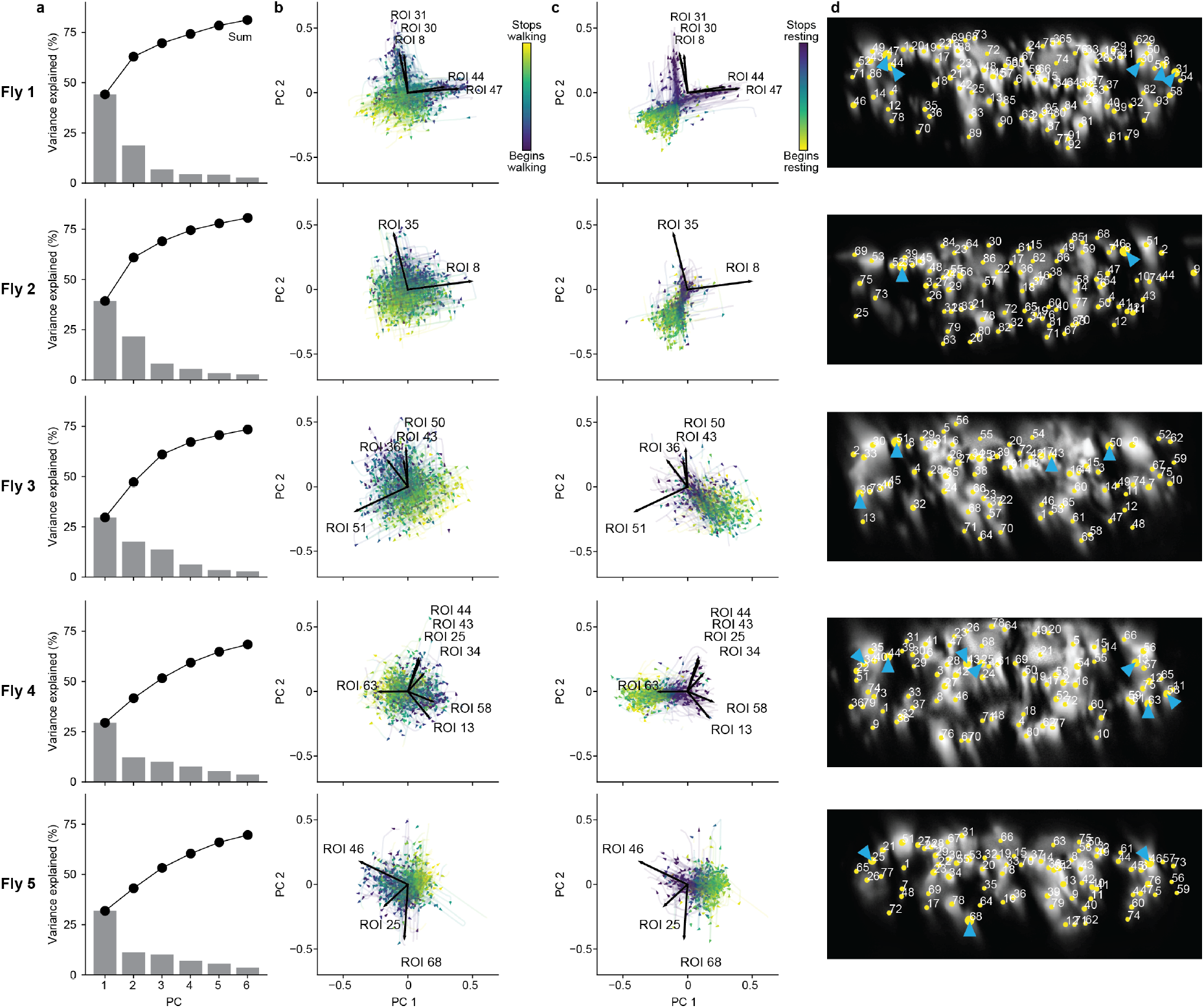
Principal component analysis of neural activity during walking and resting across individual animals. **(a)** Amount of variance explained by six principal components (PCs) of neural activity derivatives during walking for five individual flies. **(b**,**c)** The derivative of neural activity during **(b)** walking and **(c)** resting. PC embeddings were trained on data taken during walking only. Colored trajectories are individual epochs of **(b)** walking and **(c)** resting. Time is color coded and the temporal progression of each epoch is indicated (arrowheads). Note that color scales are inverted to match the color at transitions between walking and resting. Black arrows indicate PC loadings for DNs with vectors longer than 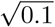. ROI numbers correspond to the connective image in panel **d. (d)** Locations of ROIs for each individual animal (yellow circles). Numbers are based on the order in **Figure S3b**. Circle radii indicate the norm of the loadings for PCs 1 and 2 (i.e., the lengths of the vectors in panels **b** and **c**). ROIs with the largest loadings are indicated (cyan arrowheads).

**Figure S6:**
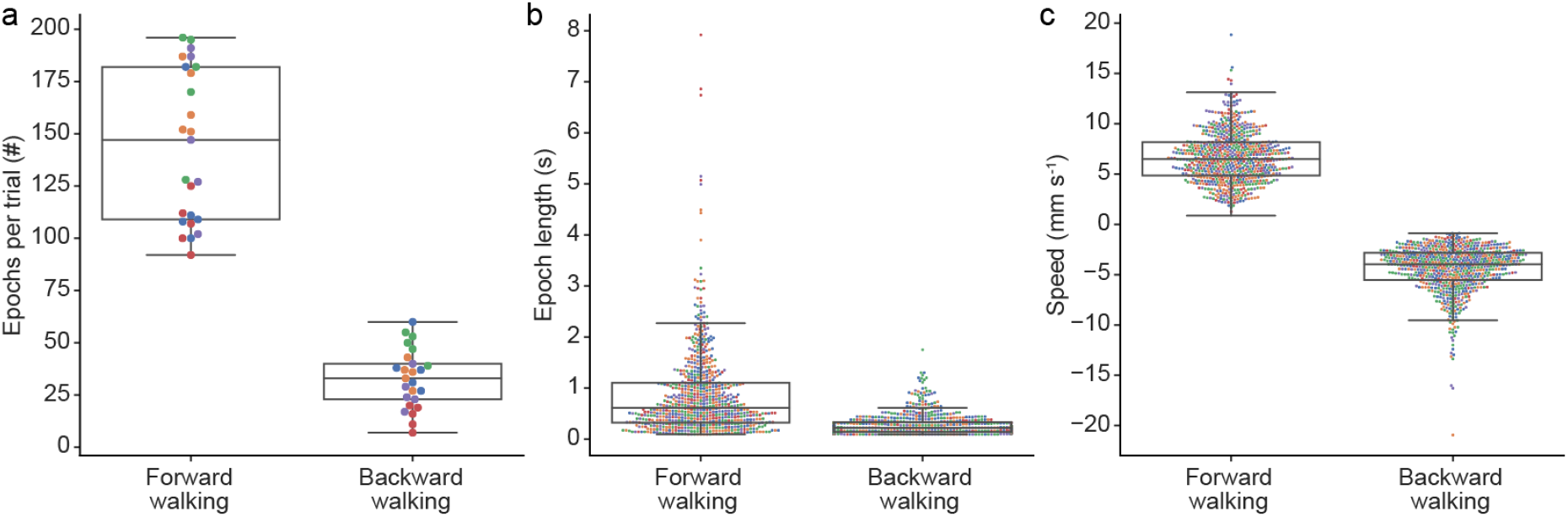
Backward walking is infrequent and brief. Compared to forward walking epochs, in our data we measured **(a)** fewer and **(b)** shorter backward walking epochs. **(c)** There was also a limited dynamic range in backward walking speed. In all panels, colors indicates fly identities. In panels **b** and **c** circles indicate the values of 700 randomly selected epochs (subsampled for clarity). Box plots indicate the median, lower, and upper quartiles. Whiskers signify 1.5 times the interquartile range.

**Figure S7:**
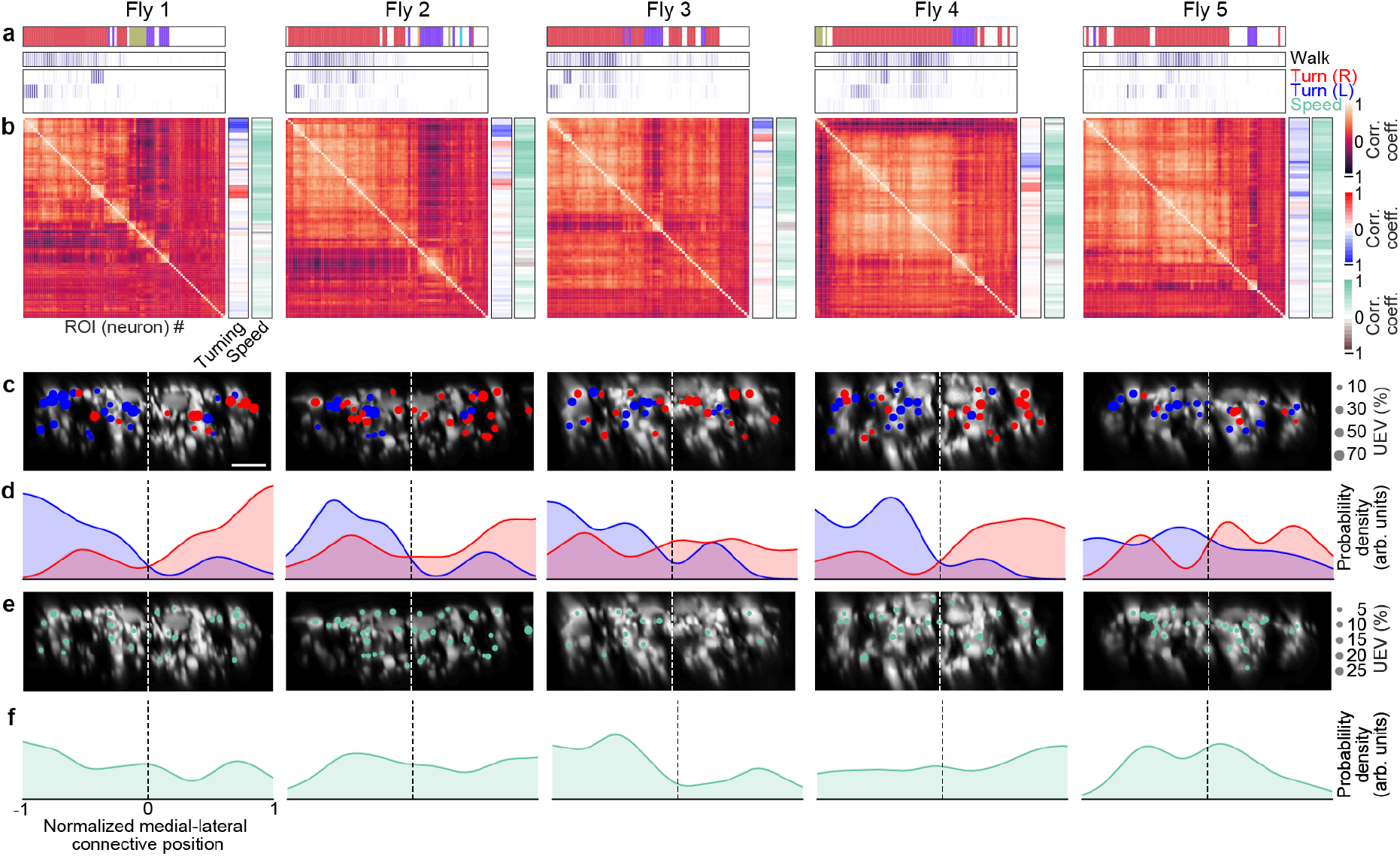
Turning and speed encoding in descending neuron populations across individual animals. **(a)** Unique explained variance of each DN from individual animals for left and right turning or walking speed. Walking *R*^2^ values are taken from **Figure S3b**. The model for turning and speed was obtained using behavior regressors as well as neural activity. To compute the UEV, activity for a given neuron was temporally shuffled. The behavior whose variance is best explained by a given neuron is indicated (top). Neurons are ordered as in panel **b. (b)** Pearson’s correlation coefficient matrix comparing neural activity across DNs is ordered by hierarchical clustering. Shown as well are the correlations of each DN’s activity with right, left, and forward walking (right). Neurons are ordered as in panel **a. (c)** Locations of turn encoding DNs (UEV *>* 5%), color-coded by preferred direction (left, blue; right, red). Circle radii scale with UEV. Dashed white line indicates the approximate midline of the cervical connective. Scale bars are 10 µm for all subpanels and panel **e. (d)** Kernel density estimates of the distributions of turn encoding DNs across individuals (opaque lines). Probability densities are normalized by the number of DNs along the connective’s medial-lateral axis. **(e)** Locations of speed encoding DNs (UEV *>* 2%). Circle radii scale with UEV. Dashed white line indicates the approximate midline of the cervical connective. **(f)** Kernel density estimates of the distributions of speed encoding DNs across individuals (opaque lines). Probability densities are normalized by the number of DNs along the connective medial-lateral axis.

**Figure S8:**
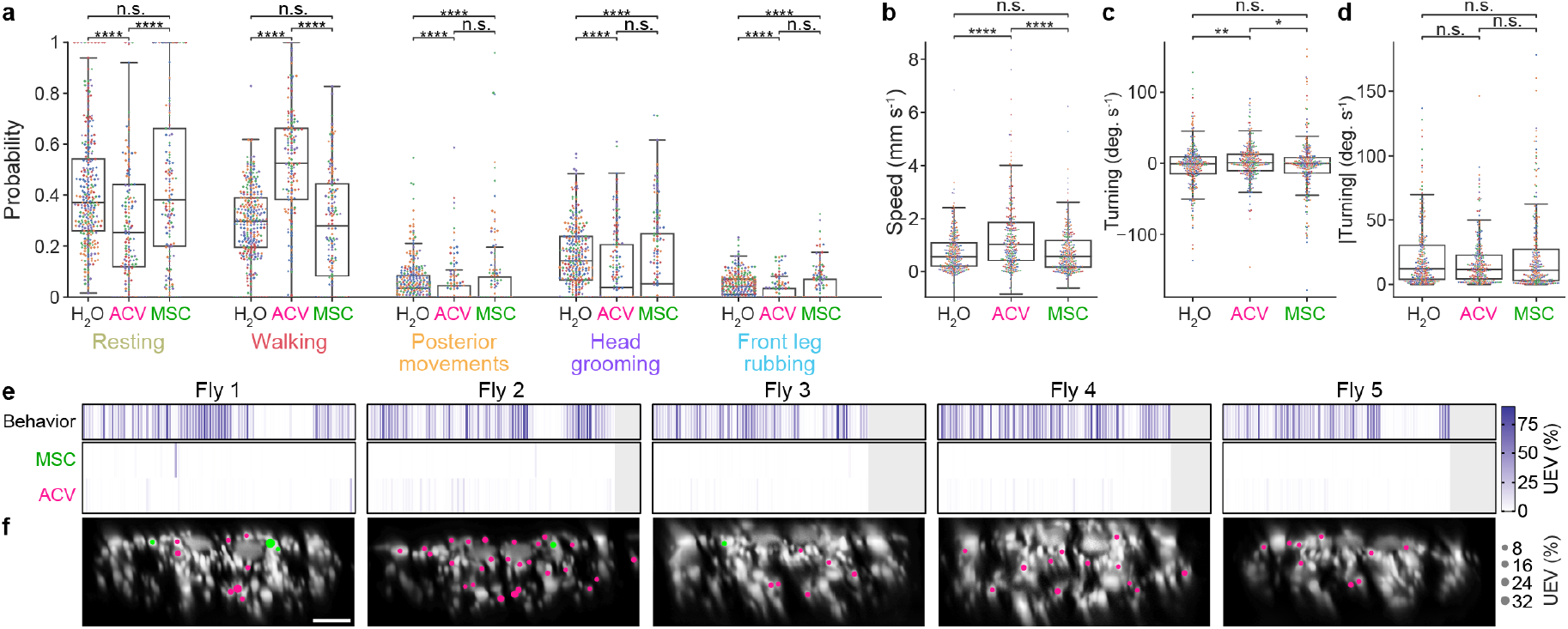
Odor-modulated behaviors and encoding in descending neuron populations across individuals. **(a)** The probabilities that classified behaviors occur during periods of stimulation with humidified air, ACV, or MSC odors. Stars indicate the significance level for a two-sided Mann-Whitney U test (‘*’: *p <* 0.05, ‘**’: *p <* 0.01, ‘****’: *p <* 10^−4^). **(b-d)** Swarm plots showing a subset of 250 randomly sampled points indicating **(b)** walking speed, **(c)** turning angular velocity, and **(d)** absolute value of turning angular velocity during periods of stimulation with humidified air, ACV, or MSC odors. **(e)** Matrices showing the cross-validated ridge regression UEV of models that contain behavior and odor regressors for five individual animals. The first row (‘Behavior’) shows the composite *R*^2^ for all behavior regressors and shuffled odor regressors. The second and third rows show UEVs for the odor regressors MSC and ACV, respectively. **(f)** Locations of odor encoding neurons (UEV *>* 5%) across five individual animals. Scale bar is 10 µm and applies to all images. Color indicates odor. Radii scale with the amount of variance explained.

**Figure S9:**
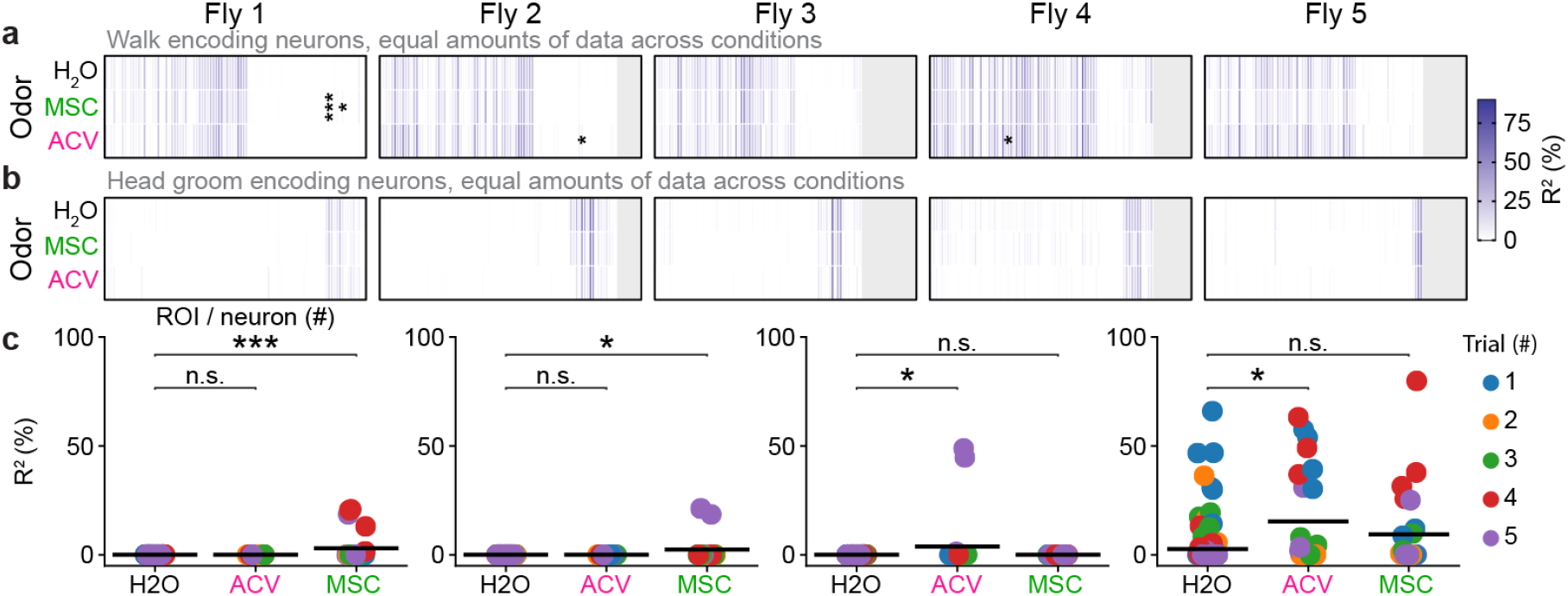
Largely identical descending neuron populations are recruited during walking and head grooming irrespective of odor context. **(a-b)** Amount of **(a)** walking or **(b)** head grooming variance explained by each DN using only frames during presentation of humidified air, apple cider vinegar (ACV), or methyl salicylate (MSC). Data during humidified air presentation were split into groups to match the amount of data available during ACV and MSC presentation. Indicated are cases where a two-sided Mann-Whitney U test comparing the cross-validation folds between humidified air and each of the odors yielded significant differences after Bonferroni correction for multiple comparisons (*: *p <* 0.05; ***: *p <* 0.001). **(c)** Individual ROI data points resulting in significant differences across conditions by a two-sided Mann-Whitney U test (shown from left to right: fly 1, ROI 82; fly 1, ROI 87; fly 2, ROI 74; fly 4, ROI 28). Each point is the result of a single cross-validation fold from a single trial.

## 8 Supplementary Videos

**Video 1: Representative recording and processing of descending neuron population activity and animal behavior**.

Fly behavior as seen by camera 5 **(top left)**. Odor stimulus presentation is indicated **(top right)**. Fictive walking trajectory of the fly calculated using FicTrac **(middle left)**. Trajectory turns gray for points after 2 s. 3D pose of the fly calculated using DeepFly3D **(bottom left)**. Text indicates the current behavior class (top right). Raw two-photon microscope image after center-of-mass alignment (**(top right)**. Δ*F/F* image after motion correction and denoising of the green channel **(middle right)**. Linear discriminant analysis-based low-dimensional representation of the neural data **(bottom right)**. Each dimension is a linear combination of neurons. The dimensions are chosen such that frames associated with different behaviors are maximally separated. https://www.dropbox.com/s/vx2zd2tt7vg0bym/Video1-Summary.mov?dl=0

**Video 2: Behavior-triggered average** Δ*F/F* **during walking epochs**.

Averaged Δ*F/F* images aligned with respect to behavior onset for all walking epochs. Red circles (top left in each imaging panel) indicate the onset of behavior. When the red circle becomes cyan, less then seven behavior epochs remain and the final image with more than eight epochs is shown. Shown as well is an example behavior epoch **(top left)** indicating the time with respect to the onset of behavior and synchronized with Δ*F/F* panels. https://www.dropbox.com/s/253ondwagchipfk/Video2_Walking.mov?dl=0

**Video 3: Behavior-triggered average** Δ*F/F* **during resting epochs**.

Averaged Δ*F/F* images aligned with respect to behavior onset for all resting epochs. Red circles (top left in each imaging panel) indicate the onset of behavior. When the red circle becomes cyan, less then seven behavior epochs remain and the final image with more than eight epochs is shown. Shown as well is an example behavior epoch **(top left)** indicating the time with respect to the onset of behavior and synchronized with Δ*F/F* panels. https://www.dropbox.com/s/gklmy23f8pj20m8/Video3_Resting.mov?dl=0

**Video 4: Behavior-triggered average** Δ*F/F* **during head grooming epochs**.

Averaged Δ*F/F* images aligned with respect to behavior onset for all head grooming epochs. Red circles (top left in each imaging panel) indicate the onset of behavior. When the red circle becomes cyan, less then seven behavior epochs remain and the final image with more than eight epochs is shown. Shown as well is an example behavior epoch **(top left)** indicating the time with respect to the onset of behavior and synchronized with Δ*F/F* panels. https://www.dropbox.com/s/quo2889b6im1z8w/Video4_HeadGroom.mov?dl=0

**Video 5: Asymmetric activity in a pair of DNx01s is associated with asymmetric leg-antennal collisions during antennal grooming**.

Three head grooming epochs in the same animal having leg contact with **(top)** primarily the left antenna, **(middle)** both antennae, or **(bottom)** primarily the right antenna. Shown are corresponding **(right)** neural activity Δ*F/F* images (note putative DNx01s in dashed white circles), **(middle)** behavior videos (camera 2), and **(left)** leg-antennal collisions (green) during kinematic replay of 3D poses in the NeuroMechFly physics simulation. Video playback is 0.25x real-time. https://www.dropbox.com/s/5l72w86njxjhoxx/Video5-AsymmetricGrooming.mov?dl=0

## 9 Data and code availability

Data are available at:

https://dataverse.harvard.edu/dataverse/DNs

Analysis code is available at:

https://github.com/NeLy-EPFL/DN_population_analysis

## 10 Funding

PR acknowledges support from an SNSF Project Grant (175667) and an SNSF Eccellenza Grant (181239). FA acknowledges support from a Boehringer Ingelheim Fonds PhD stipend.

## 11 Acknowledgments

We thank K. Asahina (Salk Institute, San Diego, USA) and B. McCabe (EPFL, Lausanne, Switzerland) for transgenic *Drosophila* strains. We thank J. Phelps for discussions and assistance with manual annotation of the EM dataset.

## 12 Author Contributions

F.A. - Methodology, Software, Formal Analysis, Investigation, Data Acquisition, Data Curation, Writing – Original Draft Preparation, Writing - Review & Editing.

C.L.C. - Investigation, Data Acquisition, Data Curation, Writing - Review & Editing.

P.R. - Conceptualization, Methodology, Resources, Writing – Original Draft Preparation, Writing - Review & Editing, Supervision, Project Administration, Funding Acquisition.

## 13 Competing interests

The authors declare that no competing interests exist.

## Notes

https://www.dropbox.com/s/vx2zd2tt7vg0bym/Video1-Summary.mov?dl=0

https://www.dropbox.com/s/253ondwagchipfk/Video2_Walking.mov?dl=0

https://www.dropbox.com/s/gklmy23f8pj20m8/Video3_Resting.mov?dl=0

https://www.dropbox.com/s/quo2889b6im1z8w/Video4_HeadGroom.mov?dl=0

https://www.dropbox.com/s/5l72w86njxjhoxx/Video5-AsymmetricGrooming.mov?dl=0

### Competing Interest Statement

The authors have declared no competing interest.

## References

[1] Cisek, P. & Kalaska, J. F. Neural mechanisms for interacting with a world full of action choices. Annual review of neuroscience 33, 269–298 (2010).

[2] Bouvier, J. et al. Descending command neurons in the brainstem that halt locomotion. Cell 163, 1191–1203 (2015).

[3] Capelli, P., Pivetta, C., Soledad Esposito, M. & Arber, S. Locomotor speed control circuits in the caudal brainstem. Nature 551, 373–377 (2017).

[4] Caggiano, V. et al. Midbrain circuits that set locomotor speed and gait selection. Nature 553, 455–460 (2018).

[5] Orger, M. B., Kampff, A. R., Severi, K. E., Bollmann, J. H. & Engert, F. Control of visually guided behavior by distinct populations of spinal projection neurons. Nature neuroscience 11, 327–333 (2008).

[6] Heinrich, R. Impact of descending brain neurons on the control of stridulation, walking, and flight in orthoptera. Microscopy Research and Technique 56, 292–301 (2002).

[7] Kien, J. Neuronal activity during spontaneous walking—i. starting and stopping. Comparative Biochemistry and Physiology Part A: Physiology 95, 607–621 (1990).

[8] Böhm, H. & Schildberger, K. Brain neurones involved in the control of walking in the cricket gryllus bimaculatus. Journal of Experimental Biology 166, 113–130 (1992).

[9] Olsen, S. R. & Wilson, R. I. Cracking neural circuits in a tiny brain: new approaches for understanding the neural circuitry of drosophila. Trends in neurosciences 31, 512–520 (2008).

[10] Namiki, S., Dickinson, M. H., Wong, A. M., Korff, W. & Card, G. M. The functional organization of descending sensory-motor pathways in drosophila. Elife 7, e34272 (2018).

[11] Hsu, C. T. & Bhandawat, V. Organization of descending neurons in drosophila melanogaster. Scientific Reports 6 (2016).

[12] von Reyn, C. R. et al. A spike-timing mechanism for action selection. Nature Neuroscience 17, 962–970 (2014).

[13] Schnell, B., Ros, I. G. & Dickinson, M. H. A descending neuron correlated with the rapid steering maneuvers of flying drosophila. Current Biology 27, 1200–1205 (2017).

[14] Chen, C.-L. et al. Imaging neural activity in the ventral nerve cord of behaving adult drosophila. Nature communications 9, 1–10 (2018).

[15] Ache, J. M. et al. Neural basis for looming size and velocity encoding in the drosophila giant fiber escape pathway. Current Biology 29, 1073–1081.e4 (2019).

[16] Namiki, S. et al. A population of descending neurons that regulates the flight motor of drosophila. Current Biology 32, 1189–1196.e6 (2022).

[17] Cande, J. et al. Optogenetic dissection of descending behavioral control in drosophila. Elife 7, e34275 (2018).

[18] Zacarias, R., Namiki, S., Card, G. M., Vasconcelos, M. L. & Moita, M. A. Speed dependent descending control of freezing behavior in Drosophila melanogaster. Nature Communications 9, 3697 (2018).

[19] Ache, J. M., Namiki, S., Lee, A., Branson, K. & Card, G. M. State-dependent decoupling of sensory and motor circuits underlies behavioral flexibility in drosophila. Nature Neuroscience 22, 1132–1139 (2019).

[20] Guo, L., Zhang, N. & Simpson, J. H. Descending neurons coordinate anterior grooming behavior in drosophila. Current Biology (2022).

[21] Zheng, Z. et al. A complete electron microscopy volume of the brain of adult drosophila melanogaster. Cell 174, 730–743 (2018).

[22] Phelps, J. S. et al. Reconstruction of motor control circuits in adult drosophila using automated transmission electron microscopy. Cell 184, 759–774 (2021).

[23] Lima, S. Q. & Miesenböck, G. Remote control of behavior through genetically targeted photostimulation of neurons. Cell 121, 141–152 (2005).

[24] Hampel, S., Franconville, R., Simpson, J. H. & Seeds, A. M. A neural command circuit for grooming movement control. elife 4, e08758 (2015).

[25] Bidaye, S. S., Machacek, C., Wu, Y. & Dickson, B. J. Neuronal control of drosophila walking direction. Science 344, 97–101 (2014).

[26] Bidaye, S. S. et al. Two brain pathways initiate distinct forward walking programs in drosophila. Neuron 108, 469–485 (2020).

[27] Kupfermann, I. & Weiss, K. R. The command neuron concept. Behavioral and Brain Sciences 1, 3–10 (1978).

[28] Zorović, M. & Hedwig, B. Processing of species-specific auditory patterns in the cricket brain by ascending, local, and descending neurons during standing and walking. Journal of Neurophysiology 105, 2181–2194 (2011).

[29] Rayshubskiy, A. et al. Neural circuit mechanisms for steering control in walking Drosophila. bioRxiv (2020).

[30] Mann, K., Gallen, C. L. & Clandinin, T. R. Whole-brain calcium imaging reveals an intrinsic functional network in drosophila. Current Biology 27, 2389–2396 (2017).

[31] Hermans, L. et al. Long-term imaging of the ventral nerve cord in behaving adult drosophila. bioRxiv (2021).

[32] Asahina, K. et al. Tachykinin-expressing neurons control male-specific aggressive arousal in drosophila. Cell 156, 221–235 (2014).

[33] Günel, S. et al. DeepFly3d, a deep learning-based approach for 3d limb and appendage tracking in tethered, adult drosophila. eLife 8 (2019).

[34] Moore, R. J. et al. Fictrac: A visual method for tracking spherical motion and generating fictive animal paths. Journal of Neuroscience Methods 225, 106–119 (2014).

[35] Chen, T.-W. et al. Ultrasensitive fluorescent proteins for imaging neuronal activity. Nature 499, 295–300 (2013).

[36] Sterne, G. R., Otsuna, H., Dickson, B. J. & Scott, K. Classification and genetic targeting of cell types in the primary taste and premotor center of the adult drosophila brain. Elife 10, e71679 (2021).

[37] Lecoq, J. et al. Removing independent noise in systems neuroscience data using deepinterpolation. Nature Methods 18, 1401–1408 (2021).

[38] Semmelhack, J. L. & Wang, J. W. Select drosophila glomeruli mediate innate olfactory attraction and aversion. Nature 459, 218–223 (2009).

[39] Mohamed, A. A. et al. Odor mixtures of opposing valence unveil inter-glomerular crosstalk in the drosophila antennal lobe. Nature Communications 10, 1–17 (2019).

[40] Lobato-Rios, V. et al. Neuromechfly, a neuromechanical model of adult drosophila melanogaster. Nature Methods 19, 620–627 (2022).

[41] Whiteway, M. R. et al. Semi-supervised sequence modeling for improved behavioral segmentation. bioRxiv (2021).

[42] Seeds, A. M. et al. A suppression hierarchy among competing motor programs drives sequential grooming in drosophila. eLife 3 (2014).

[43] Ravbar, P., Zhang, N. & Simpson, J. H. Behavioral evidence for nested central pattern generator control of drosophila grooming. Elife 10, e71508 (2021).

[44] Mendes, C. S., Bartos, I., Akay, T., Márka, S. & Mann, R. S. Quantification of gait parameters in freely walking wild type and sensory deprived drosophila melanogaster. elife 2, e00231 (2013).

[45] Kato, S. et al. Global brain dynamics embed the motor command sequence of caenorhabditis elegans. Cell 163, 656–669 (2015).

[46] Braitenberg, V. Vehicles: Experiments in synthetic psychology (MIT press, 1986).

[47] Sen, R. et al. Moonwalker descending neurons mediate visually evoked retreat in drosophila. Current Biology 27, 766–771 (2017).

[48] Jenett, A. et al. A gal4-driver line resource for drosophila neurobiology. Cell Reports 2, 991–1001 (2012).

[49] Chen, C.-L. et al. Ascending neurons convey behavioral state to integrative sensory and action selection centers in the brain. bioRxiv (2022).

[50] Nern, A., Pfeiffer, B. D. & Rubin, G. M. Optimized tools for multicolor stochastic labeling reveal diverse stereotyped cell arrangements in the fly visual system. Proceedings of the National Academy of Sciences 112, E2967–E2976 (2015).

[51] Harris, R. M., Pfeiffer, B. D., Rubin, G. M. & Truman, J. W. Neuron hemilineages provide the functional ground plan for the drosophila ventral nervous system. elife 4, e04493 (2015).

[52] Aimon, S. et al. Fast near-whole–brain imaging in adult drosophila during responses to stimuli and behavior. PLoS biology 17, e2006732 (2019).

[53] Schaffer, E. S. et al. Flygenvectors: the spatial and temporal structure of neural activity across the fly brain. bioRxiv (2021).

[54] Brezovec, L. E., Berger, A. B., Druckmann, S. & Clandinin, T. R. Mapping the neural dynamics of locomotion across the drosophila brain. bioRxiv (2022).

[55] Tanaka, R. & Clark, D. A. Neural mechanisms to exploit positional geometry for collision avoidance. Current Biology (2022).

[56] Triphan, T., Poeck, B., Neuser, K. & Strauss, R. Visual targeting of motor actions in climbing drosophila. Current biology 20, 663–668 (2010).

[57] Coen, P. et al. Dynamic sensory cues shape song structure in drosophila. Nature 507, 233–237 (2014).

[58] Nässel, D. R., Högmo, O. & Hallberg, E. Antennal receptors in the blowfly Calliphora erythrocephala. i. the gigantic central projection of the pedicellar campaniform sensillum. Journal of Morphology 180, 159–169 (1984).

[59] King, D. G. & Wyman, R. J. Anatomy of the giant fibre pathway in Drosophila. i. three thoracic components of the pathway. Journal of Neurocytology 9, 753–770 (1980).

[60] Azevedo, A. W. et al. A size principle for recruitment of drosophila leg motor neurons. Elife 9, e56754 (2020).

[61] Zhang, Y. et al. jgcamp8 fast genetically encoded calcium indicators (2020).

[62] Piatkevich, K. D. et al. A robotic multidimensional directed evolution approach applied to fluorescent voltage reporters. Nature Chemical Biology 14, 352–360 (2018).

[63] Voleti, V. et al. Real-time volumetric microscopy of in vivo dynamics and large-scale samples with scape 2.0. Nature methods 16, 1054–1062 (2019).

[64] Aimon, S., Cheng, K. Y., Gjorgjieva, J. & Kadow, I. C. G. Walking elicits global brain activity in Drosophila. bioRxiv (2022).

[65] Mann, K., Gordon, M. & Scott, K. A pair of interneurons influences the choice between feeding and locomotion in drosophila. Neuron 79, 754–765 (2013).

[66] Tastekin, I. et al. Role of the subesophageal zone in sensorimotor control of orientation in drosophila larva. Current Biology 25, 1448–1460 (2015).

[67] Simpson, J. H. Rationally subdividing the fly nervous system with versatile expression reagents. Journal of Neurogenetics 30, 185–194 (2016).

[68] Pavlou, H. J. & Goodwin, S. F. Courtship behavior in Drosophila melanogaster : towards a ‘courtship connectome’. Current Opinion in Neurobiology 23, 76–83 (2013).

[69] Krull, A., Buchholz, T.-O. & Jug, F. Noise2void-learning denoising from single noisy images. In Proceedings of the IEEE Conference on Computer Vision and Pattern Recognition, 2129–2137 (2019).

[70] Jefferis, G. S. et al. Comprehensive maps of drosophila higher olfactory centers: Spatially segregated fruit and pheromone representation. Cell 128, 1187–1203 (2007).

[71] Saalfeld, S., Cardona, A., Hartenstein, V. & Tomančák, P. CATMAID: collaborative annotation toolkit for massive amounts of image data. Bioinformatics 25, 1984–1986 (2009).

[72] Schneider-Mizell, C. M. et al. Quantitative neuroanatomy for connectomics in drosophila. Elife 5, e12059 (2016).

[73] Bogovic, J. A. et al. An unbiased template of the drosophila brain and ventral nerve cord. Plos one 15, e0236495 (2020).

[74] Musall, S., Kaufman, M. T., Juavinett, A. L., Gluf, S. & Churchland, A. K. Single-trial neural dynamics are dominated by richly varied movements. Nature Neuroscience 22, 1677–1686 (2019).

[75] Virtanen, P. et al. SciPy 1.0: Fundamental Algorithms for Scientific Computing in Python. Nature Methods 17, 261–272 (2020).

[76] Pedregosa, F. et al. Scikit-learn: Machine learning in Python. Journal of Machine Learning Research 12, 2825–2830 (2011).

[77] Chartrand, R. Numerical differentiation of noisy, nonsmooth data. International Scholarly Research Notices 2011 (2011).

[78] Jr., J. H. W. Hierarchical grouping to optimize an objective function. Journal of the American Statistical Association 58, 236–244 (1963).

[79] Jiao, W. et al. Intact drosophila whole brain cellular quantitation reveals sexual dimorphism. bioRxiv (2021).

